# Endogenous Real Time Imaging Reveals Dynamic Chromosomal Mobility During Ligand-Mediated Transcriptional Burst Events

**DOI:** 10.1101/2025.08.18.670875

**Authors:** Susan Wang, Thomas Suter, Amir Gamliel, Yeeun Kim, Sreejith J. Nair, Soohwan Oh, Feng Yang, Kenneth A. Ohgi, Tobias Wagner, Steven Gan, Michael G. Rosenfeld

## Abstract

Enhancers serve as the major genomic elements regulating mammalian signal-dependent transcriptional programs, characterized by alternating periods of target gene “bursting” and “non-busting” that require investigation of induced enhancer condensates and locus motility in real time to provide dynamic insights into signal/ligand-dependent regulatory events. Here, endogenous live cell imaging has revealed the altered chromosomal dynamics/condensate formation occurring during estrogen receptor α (ERα)-dependent target gene bursting/post-bursting and chronic activation events. Simultaneous DNA/RNA endogenous live imaging reveals that an increased mobility of acutely ERα-stimulated loci observed during the bursting phase is, unexpectedly, further increased in the subsequent non-burst phase. Single molecule tracking (SMT) of ERα shows that the relatively high-burst, lower-mobility acute state was indeed enriched for high-viscosity, 1,6-hexanediol-sensative ERα molecules in a low sub-diffusive confined state with enhanced condensate formation during burst activation. Consistent with this, blocking transcription with flavopiridol shifts DNA tracks into a non-confined state. Differential DNA kinetics during burst vs non-burst has provided a strategy to assess altered condensate formation during gene activation events. (165)

## Main

Forty years removed from their initial discovery, gene regulation by transcriptional enhancers has been revealed to be one of the dominant molecular mechanisms underlying cell-type and signal-specific transcriptional diversity in metazoans^1–3^. In addition to their well-documented critical functions in cell type determination and differentiation, transcriptional enhancers have proved to serve as the predominant regulators of the precise, rapidly altered patterns of gene regulation in response to diverse acute signaling pathways, reflecting the preferential binding of the majority of these signal-dependent transcription factors to enhancers, rather than to promoters^4–11^. Indeed, signal-dependent enhancers themselves are active transcription units producing noncoding enhancer RNAs (eRNAs) with concomitant condensation of RNA-dependent ribonucleoprotein (RNP) complexes upon acute stimulation^12–19^, with their activation generally temporally preceding activation of its cognate promoter.

The cell nucleus has long been recognized to harbor sub-nuclear architectural structures, including the nucleolus, nuclear speckles, paraspeckles, and Cajal bodies, several of which are assembled as biomolecular condensates proposed to involve liquid–liquid phase separation (LLPS)^20–24^. In addition to the complex recruitment events occurring at activated enhancers, it has been recognized that the potential interactions of enhancers with subnuclear architectural structures mediate transcriptional activation. Actively transcribed, enhancer-rich genomic regions are segregated near interchromatin granules while the majority of the transcriptionally inactive regions are associated with nucleoli^25–35^. A poorly characterized nuclear assembly built around matrin-3 protein, a DNA– RNA binding protein with diverse biological function, is another such non-membrane-bound nuclear compartment implicated to contribute to transcription regulation upon colocalization with target enhancers^36,37^. A key insight into the regulation of transcription has been the understanding that transcription occurs in bursts of high transcriptional activity followed by non-burst periods of low transcription, although the relative intensity, duration, and frequency of bursts have been shown to vary between genes and signaling^38–40^.

Therefore, it now becomes critical to integrate the spatial and temporal aspects of signal-induced enhancer activation and alterations in the putative enhancer condensates, motility of the chromosomal loci and their relationship to subnuclear architectural structures. A particularly instructive model has been provided by the regulation of a large program of acute ligand-dependent enhancer activation in breast cancer cells (MCF7), causing the rapid assembly of RNP complexes, referred to as the MegaTrans complex, at the ∼1-2000 most robustly activated ERα-bound enhancers with RNP condensate-like properties controlling E_2_-regulated transcriptional programs^25,41^. In breast cancer cells, estradiol-17β stimulation results in the activation of hundreds of non-clustered enhancers with very high levels of eRNA transcription and higher recruitment of chromatin regulators^42–45^. The MegaTrans complex is mainly composed of other TFs such as GATA3, AP2γ, RARα, YAP/TEAD, and non-classical cofactors that are recruited to the chromatin indirectly, bringing together a large ribonucleoprotein complex as a result of high-affinity chromatin binding of a single TF^41,46–49^.

In light of our labs’ previous findings of distinct expression patterns in acute and chronic stimulation of breast cancers cells by ERα signaling, we sought to explore whether these two states of ERα signaling operated under different modes of nuclear architectural dynamics, perhaps reflecting differential MegaTrans enhancer condensate properties. Live-imaging approaches that combined endogenous spatiotemporal visualization of ERα target-gene and transcript identified distinct DNA and RNA signatures for both the acute and chronic stimulation phases and allowed us to identify a novel chromatin kinetic signature associated with transcriptional bursting. These data revealed a multi-state model of chromatin kinetics in transcriptional activation: a low mobility, pre-stimulation state; a high-mobility, non-burst stimulated state; and an intermediate-mobility, burst-stimulated state. Interestingly, we observed a high-viscosity, 1,6-hexanediol sensitive subset of ERα molecules that is enriched in the intermediate-mobility, burst-stimulated state. In the absence of transcriptional bursting, that ERα fraction along with chromatin are particularly unrestricted and increases in motility, suggesting confinement via condensate formation during transcriptional bursting events.

## Results

### Acute vs chronic E_2_ enhancer activation-dependent bursting

To investigate the dynamic acute vs. chronic regulation of estradiol-17β-dependent enhancer-activated gene bursting, we selected the *NRIP1* locus for tagging, based upon its well-established ERα enhancer responsivity, its high level of transcription, and its diploid status in MCF7 facilitates DNA-labeling of the same allele as the RNA tagged transcription unit. We incorporated a 24x MS2 cassette into the 3’ UTR of the *NRIP1* target gene, which forms a series of MS2-hairpin loops upon transcription that can rapidly bind MS2-coat-protein (MCP) fluorescent-protein fusions with high specificity and affinity, resulting in punctate fluorescent signal coincident with *NRIP1* gene transcription^50–53^. MS2 insertion into the endogenous loci was facilitated by CRISPR-Cas9, and selection was improved through the addition of either a fluorescent or puromycin selection marker to the homologous recombination (HR) donor cassette just upstream of the stop codon **(Fig. 1a).** MCP-YFP fusion proteins were then stably integrated into these lines via lentiviral infection, and individual clones were screened for the ability to show transient punctate foci in response to E_2_ treatment **(Fig. 1b; Supplemental Movie 1)**. Accurate marking in the selected clones were validated by FISH, with the line used for further analyses showing 94.7% overlap between punctate YFP signal and *NRIP1* gene FISH signal **(Fig. 1c).**

**Figure 1.**
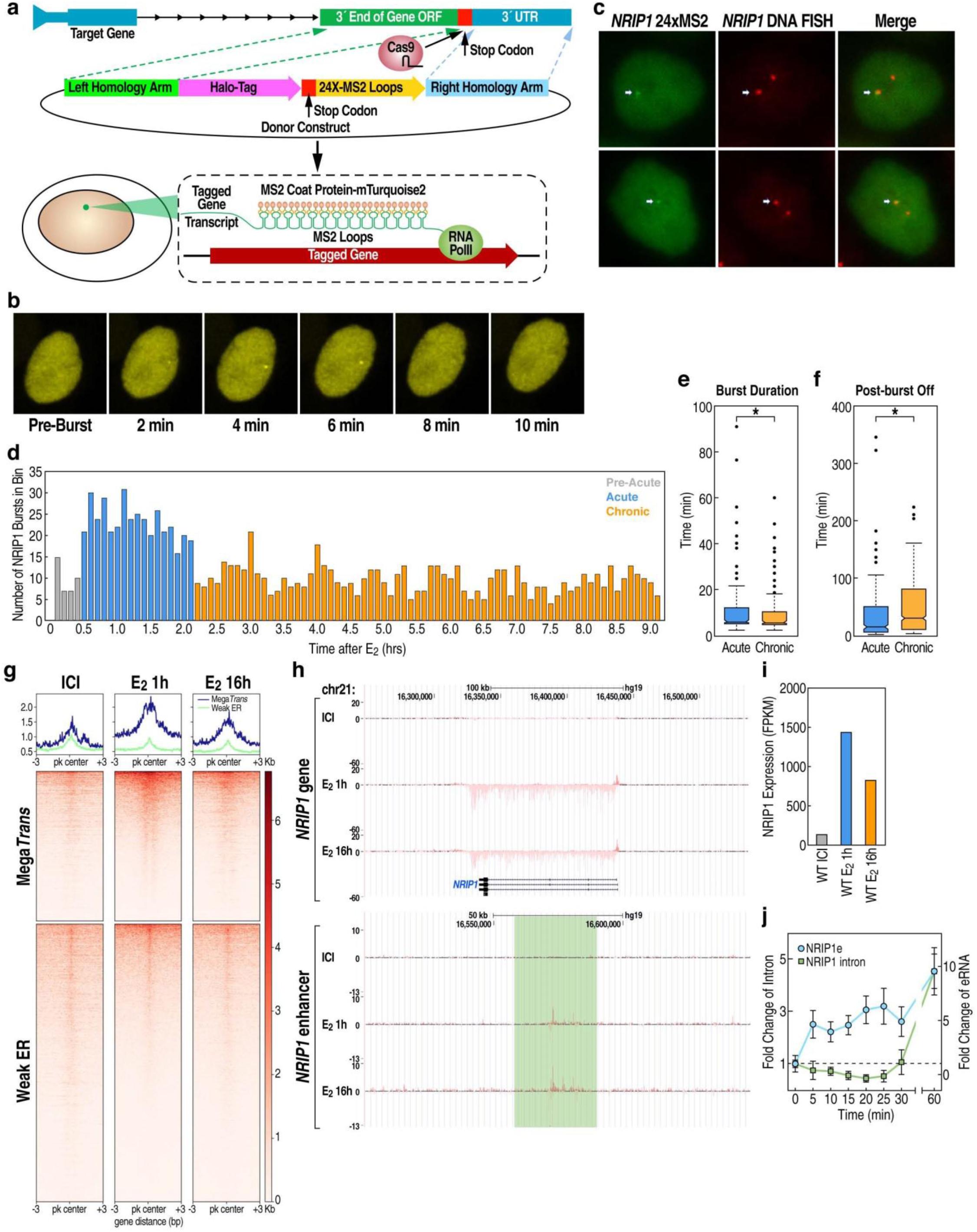
RNA visualization of transcriptional bursting in acute and chronically ERα stimulated states. **a.** RNA visualization achieved via 24xMS2 insertion into the endogenous NRIP1 loci via CRISPR-Cas9, and selection was improved through the addition of either a fluorescent or puromycin selection marker to the HR donor cassette just upstream of the stop codon. **b.** An example of a bust event at NRIP1 loci in response to E_2_ stimulation. **c.** RNA visualization FISH validation of tagged NRIP1 allele using 24xMS2 showed 94.7% overlap (n=18). **d.** Long-term timelapse show 3 clear phases of bursting in response to E_2_ treatment: high bursting post-E_2_ 3 clear phases of bursting in response to E_2_ treatment: high bursting activity from 24 to 132 minutes post-E_2_ treatment (Acute); sustained lower bursting activity immediately following Acute phase (Chronic); and low bursting activity post-E_2_ but preceding Acute phase (pre-Acute). (n=922) **e.** Acute and chronic bursts of NRIP1 differ significantly in duration, with longer on burst duration during acute phase. **f.** Acute and chronic bursts of NRIP1 differ significantly in frequency, with longer off-periods in chronic phase. **g.** PRO-seq of Mega-Trans and ER bound loci under minus, acute, and chronic E_2_ induction. **h.** GRO-seq of NRIP1 enhancer and gene expression levels shown on genome browser. **i.** GRO-seq showing NRIP1 expression levels (FPKM) are ∼1.8x higher in the acute relative to the chronic state. **j.** Time-course of NRIP1 enhancer and gene transcript levels after E_2_ stimulation measured by qPCR. (All statistics: 2-Tailed Paired T Test * = p<0.05, ** = p<0.01, *** = p<0.001, **** = p<0.0001)

To quantify burst kinetics under both acute and chronic E_2_-activated conditions, we treated cells with E_2_ and imaged at 2-minute intervals for 9 hours (minus E_2_ condition showed negligible bursting). Interestingly, we observe 3 clear phases of bursting in response to E_2_ treatment: a period of high bursting activity from 24 to 132 minutes post-initial E_2_ treatment, referred to as the acute activation period; a period immediately following the acute with bursting activity approximately half that of the acute rate, referred to as the chronic activation period; and a period between 0 and 24 minutes post-E_2_ treatment, referred to as the basal/pre-acute activation period **(Fig. 1d).** We observed that acute and chronic bursts differed significantly in both their duration and frequency **(Fig. 1e, 1f),** consistent with ∼40% diminished target gene transcriptional activation in the chronic phase observed by PRO-seq for both *NRIP1* specifically as well as MegaTrans and ER-bound regulatory enhancers **(Fig. 1g-1i)**. qPCR further validated that like most genes, *NRIP1* E_2_-dependent enhancer activation precedes activation of cognate promoter, suggesting a fundamental alteration of enhancer-regulated transcriptional mechanisms between acute and chronic E_2_ signaling **(Fig. 1j).**

### Mobility of the NRIP1 locus increases with chronic vs acute E_2_-dependent activation

We next established a system for the live imaging of the NRIP1 loci to determine a locus’s relative mobility in the acute versus chronic states, based on tagging the locus by integration of a 144xCuO cassette 2.5Kb upstream of the *NRIP1* promoter. Given the inefficiencies of homologous recombination in insertions of long repeat cassettes, which include low integration efficiency and reduction of repeat number, we first introduced a 65bp integrase site into the loci via Cas9^54^. The 144xCuO array was then integrated into the site via introduction of an Bxb1 integrase expression plasmid along with a repeat donor cassette containing a Bxb1 attB integrase target site **(Fig. 2a).** The CymR protein, which binds to the CuO repeats with high affinity and specificity, was fused to either mTurquoise2, mKate2, or Halo and stably expressed in the CuO repeat-containing cells **(Supplemental Fig. 1; Supplemental Movie 2)**. The resultant fluorescent puncta were validated as corresponding to *NRIP1* via DNA FISH, exhibiting 100% overlap **(Fig. 2b).**

**Figure 2.**
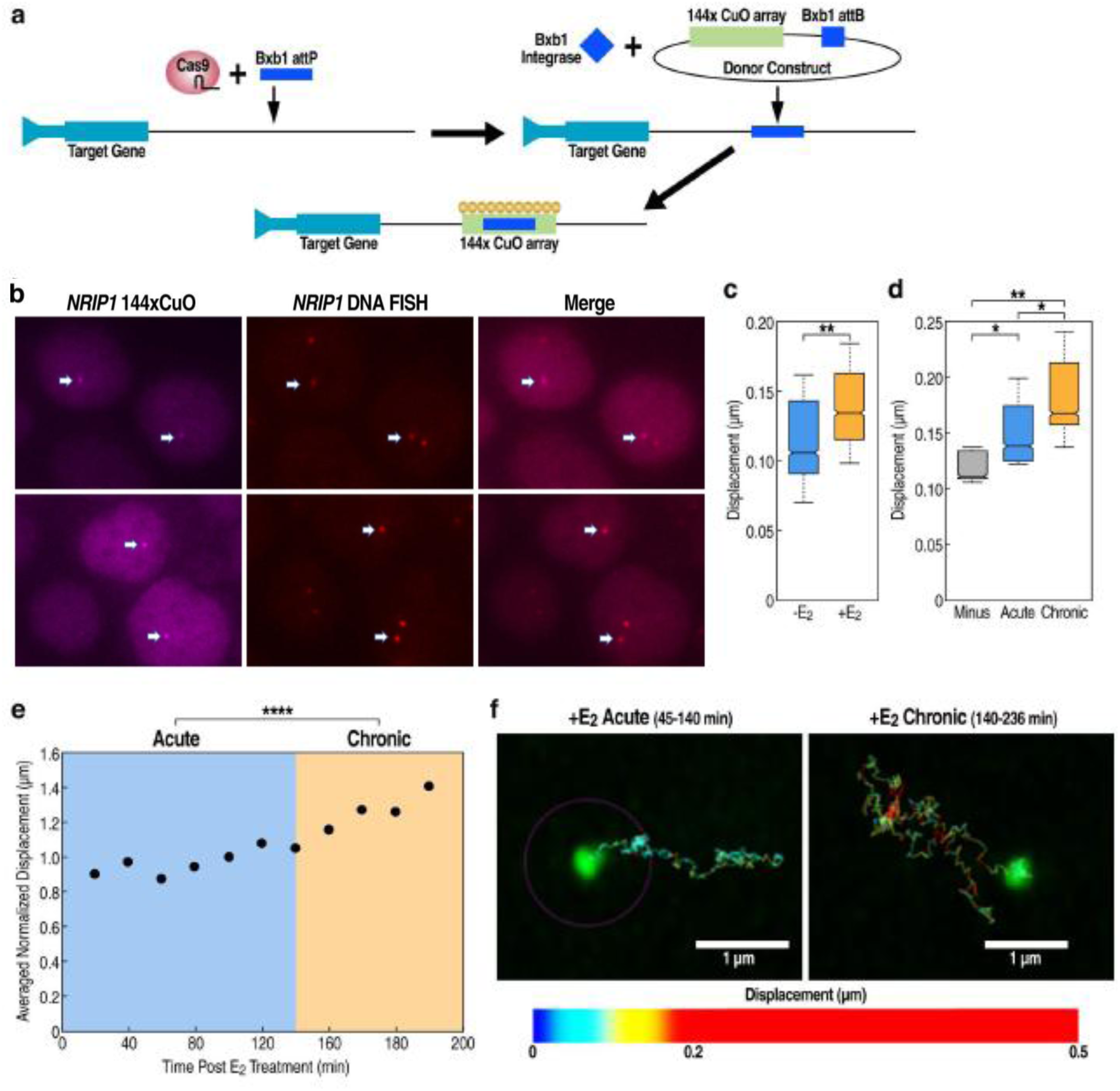
DNA visualization of loci kinetics in acute versus chronic ERα induced states. **a.** DNA visualization via endogenous tagging of *NRIP1* by knock-in of 144xCuO using Bxb1 integrase. **b.** DNA visualization FISH validation of tagged *NRIP1* allele using 144xCuO showed 100% overlap. **c.** E_2_ minus condition shows significantly lower mobility than stimulated E_2_ state (n=15). **d.** E_2_ minus condition shows significantly lower mobility than either the acute or chronic stimulation states, but there is also a significant further increase in mobility in the chronic versus acute stimulation states. **e.** Time-course of averaged normalized displacement of *NRIP1* under acute versus chronic stimulation. **f.** Contrails of *NRIP1* loci in acute versus chronic states, showing increased displacement under chronic E_2_ stimulation. (All statistics: 2-Tailed Paired T Test * = p<0.05, ** = p<0.01, *** = p<0.001, **** = p<0.0001)

We then assessed the mobility of the *NRIP1* locus by comparing the change in its average displacement per frame between acute, chronic, or no E_2_ stimulation conditions. Consistent with a previous report that loci in more transcriptionally active states showed higher mobility^55^, we noted that the E_2_-minus condition showed significantly lower mobility than either the acute or chronic stimulation states **(Fig. 2c)**. However, rather surprisingly, we detected a significant further <u>increase</u> in mobility with chronic (versus acute) E_2_ stimulation, in contrast with higher transcriptional activation of the *NRIP1* enhancer and target in the acutely-activated state **(Fig. 2d, 2e).** DNA contrails of *NRIP1* loci further demonstrated an increased displacement in chronic state compared to acute **(Fig. 2f).** Thus, while our mobility data confirms the association of transcriptional activation with increased chromosomal mobility in certain activation contexts, in line with published literature, these results suggest a necessary distinction between the initial E_2_-induced enhancer condensate, imposing reduced motility, compared to the subsequent altered condensates following chronic E_2_ treatment, licensing increased motility.

### Dynamics of ligand-dependent alterations in interactions with estrogen receptor

To better understand the mechanisms underlying the differential mobility observed for ERα stimulated loci, we further investigated the dynamic interactions of ERα in minus, acute, and chronic states. Stimulation with E_2_ results in the formation of clear ERα foci, and we observe that these foci are distinct from, but in close proximity to, both the SC35 and the Matrin3-rech nuclear network **(Fig. 3a).** We also detect a substantial increase in ERα and MegaTrans binding genome wide upon E_2_ induction via ChIP-seq, where binding is twice as robust during the acute stimulated phase relative to the chronic **(Fig. 3b, 3c).** Therefore, we sought to assess whether the biophysical properties or distribution of ERα itself are altered in acute versus chronic ERα stimulation.

**Figure 3.**
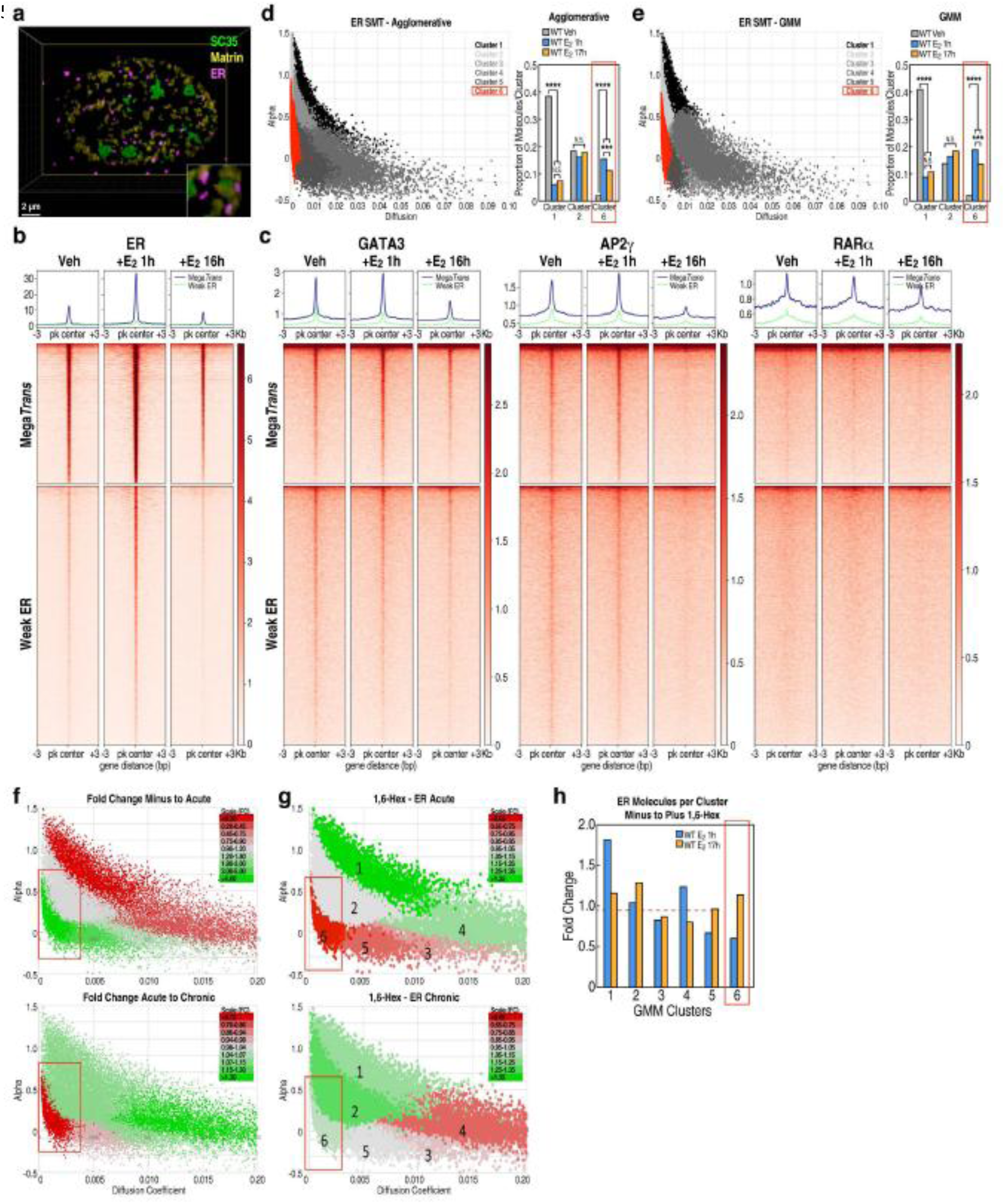
Biophysical insight into transcription dynamics via single molecule tracking. **a.** Live imaging shows that ERα foci are distinct from but in close proximity to both SC35 and the nuclear matrix. **b.** ERα ChIP-seq shows increased ERα binding genome wide upon E_2_ induction, where ER is more robustly bound in the acute phase relative to chronic. **c.** ChIP-seq of MegaTrans factors GATA3, AP2γ, and RARα show similar increased genome wide binding upon E_2_ induction, with more robust binding in acute versus chronic states. **d.** Agglomerative clustering shows an acute specific population of molecules (cluster 6, red) with the lowest diffusion coefficient and varies in sub-diffusivity as reflected by the range in alpha. (n=230) **e.** GMM shows the same acute specific population of molecules (cluster 6, red) with the lowest diffusion coefficient and varies in sub-diffusivity as reflected by the range in alpha. **f.** Significant enrichment of the lowest diffusion-coefficient, low-subdiffusivity fraction in the acute versus chronic stimulations. **g.** Analysis of 1,6-hexandiol treatment shows particular disruption of ERα molecules from acute specific cluster 6. **h.** Bar graphs show fold change of ERα molecules in each GMM cluster when treated with 1,6-hexandiol, with cluster 6 showing the greatest difference in fold change between acute and chronic states. (All statistics: 2-Tailed T Test * = p<0.05, ** = p<0.01, *** = p<0.001, **** = p<0.0001)

We performed high temporal resolution (5ms) SMT on ERα to determine the acute and/or chronic sensitivity of any ERα populations, and whether the acute and/or chronic differential mobility of these fractions corresponds with the differential chromosomal mobility observed in chronic ERα stimulation. To permit execution of SMT of ERα at high temporal resolutions, we integrated a Halo-tag into the 3’ end of the endogenous *ESR1* gene in MCF7 cells to permit stable Halo-tagging of ERα expressed at endogenous levels. Halo-tagged ERα molecules were then fluorescently labeled the Janelia Fluor 549 and imaged cells at basal conditions or after acute or chronic E_2_-induced ERα stimulation at 5ms resolution using HILO microscopy **(Supplemental Movie 3-5).**

For each ERα SMT track, six properties were calculated: 1) diffusion coefficient, 2) sub-diffusivity (alpha), 3) radial diffusion coefficient, 4) radius of confinement, 5) average 5ms displacement, and 6) duration of binding in a 250nm radius. Tracks were then clustered based upon these six parameters using either Gaussian Mixture Models (GMM) or Agglomerative Hierarchical clustering. Our clusters show the expected differences between the no-stimulation and the acute or chronic E_2_-induced stimulation. In low-diffusive states, no stimulation shows enrichment of ER in the high alpha unbound fraction, while the stimulation results in clear enrichment in the low alpha sub-diffusive restricted fractions **(Fig. 3d, 3e; Supplemental Fig. 2a, 2b)**. However, we see a highly significant enrichment of the lowest diffusion-coefficient, low sub-diffusivity fraction in the acute stimulation, whereas the higher diffusion coefficient, low sub-diffusivity fractions were unchanged or enriched in the chronic stimulation condition **(Fig. 3f).** These results suggest the acute stimulation is enriched for a very low diffusion-coefficient, restricted and potentially higher density population, which makes up roughly 8% of all ERα single molecules. That fraction dissipates into higher diffusion-coefficient populations with extended, chronic stimulation. Further insight is provided by the observation that treatment of cells with 1,6-hexanediol results in the disruption of that low diffusion-coefficient and potentially higher density population of ERα in acutely, but not in chronically treated, cells **(Fig. 3g, 3h),** which is consistent with previous observations that activation of MegaTrans enhancers is disrupted by 1,6-hexanediol^25^. Taken together, our ERα SMT data suggests the presence of a relatively highly viscous and potentially phase-separated condensate upon E_2_ stimulation and implicates these, and possibly additional, condensates in the low-mobility dynamics of the acutely-stimulated condition.

### Distinct motility states of E_2_-activated loci during bursting and non-bursting periods

Given the unexpected relationship between burst frequency and DNA mobility in the acute and chronic states as well as the identification of an acute-specific, low sub-diffusion high density ERα fraction, it was important to directly compare RNA bursting with DNA mobility at the regulated locus during the transition from bursting to non-bursting intervals. Because analyses that lack measurement of direct transcriptional output preclude determination of whether any increased DNA mobility is associated with the burst or the non-burst phases at a stimulated locus, it was important to determine whether these two phases exhibit distinct, and previously undetermined, kinetic properties that could account for the acute versus chronic dynamics, such as those observed in Figure 2. To permit resolution of this question, we engineered a cell line that permitted a combination of RNA-burst visualization and DNA-visualization methodologies in the same cell **(Fig. 4a).** We first confirmed via imaging that both the DNA and RNA tags were on the same allele **(Fig. 4b, 5e).** With this validation, we were able to observe a significant and dramatic reduction in mobility that temporally corresponds to the period of gene bursting **(Fig. 4c, 4d)**. These results highly suggest that the burst and non-burst phases of stimulated loci exist in two distinct mobility states, consistent with models of transcription that involve formation of a co-factor rich, high-valency condensate, as previously documented for several acutely-induced enhancer activation programs.

**Figure 4.**
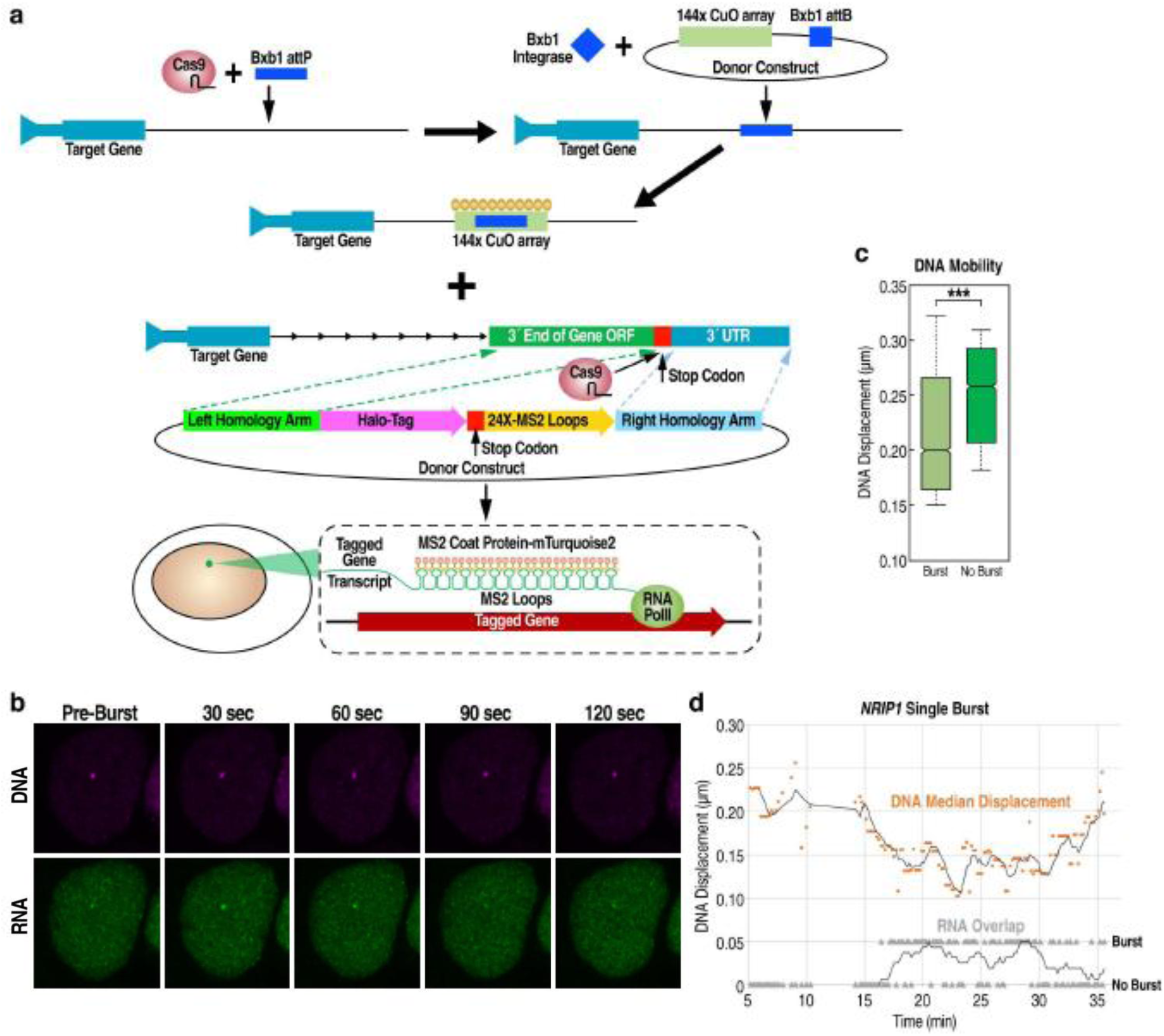
DNA RNA dual visualization of mobility relative to transcriptional bursting. **a.** Addition of RNA-burst visualization into DNA-visualization NRIP cell line. **b.** DNA RNA overlap in live cells confirms tagging of same NRIP allele. **c.** DNA mobility corresponds to gene bursting, where a significant reduction in mobility occurs during burst. (n=10) **d.** Graph mapping DNA displacement against an RNA burst, showing decrease in displacement during a burst event (characterized in binary form of 0 for no bust and 0.05 for burst signal, trend line is a rolling average of burst signal). (All statistics: 2-Tailed Paired T Test * = p<0.05, ** = p<0.01, *** = p<0.001, **** = p<0.0001)

### Burst events in proximity to subnuclear architectural structures

The potential functional importance of increased proximity of a regulated gene locus to specific subnuclear architectural structures that may facilitate condensate formation during burst activation has been of particular interest. The nucleoli, PML bodies, Matrin3 network, and the polycomb complex have also been reported to have functional impacts on nuclear organization and dynamics, with interchromatin granules and Matrin3 network both implicated in ERα-regulated transcription^25,37^. Therefore, to better understand the mechanisms underlying these associations, we investigated the dynamics of *NRIP1* relative to interchromatin granules and the Matrin3 network. We found a clear E_2_-induced increase in proximity of the *NRIP1* loci with both the Matrin3-rich network and the SC35 (representative of interchromatin granules) via immuno-DNA FISH, with the interchromatin granule’s proximity to *NRIP1* peaking in the acutely-stimulated condition while the Matrin3-rich network’s proximity being enriched in the chronic E_2_ condition **(Fig. 5a, 5b).** We have reproduced this same finding in a separate ERα-regulated gene, *TFF1* **(Supplemental Fig. 3a, 3b).**

**Figure 5.**
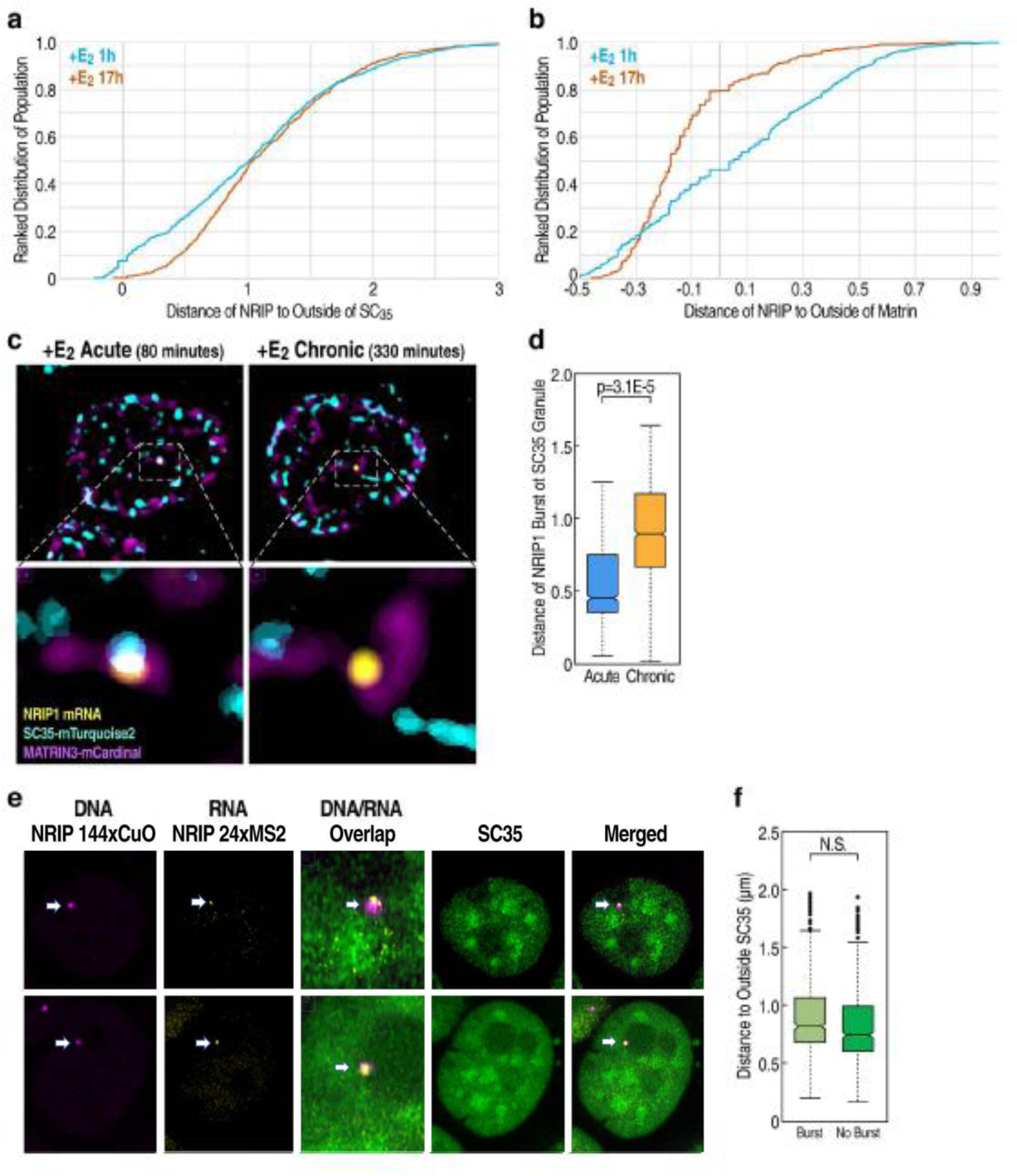
ERα regulated locus and its association with nuclear sub-structures. **a.** Immuno-DNA FISH of *NRIP1* shows increased proximity to SC35 in acute phase relative to chronic. (n=822) **b.** Immuno-DNA FISH of *NRIP1* shows induced association with Matrin3 network in the chronic state relative to acute. **c.** Three-color live visualization of *NRIP1* RNA via 24xMS2 and virally transduced SC35 and Matrin3 fused to fluorescent tags. **d.** Live imaging of *NRIP1* relative to SC35 shows peak induced proximity during the acute phase versus chronic. **e.** Three-color living visualization of *NRIP1* DNA, *NRIP1* RNA, and endogenously tagged SC35. **f.** Although E_2_ stimulation caused induced proximity of *NRIP1* to SC35 in general, there is no correlation between proximity and a burst event. (n=10) (All statistics: 2-Tailed Paired T Test * = p<0.05, ** = p<0.01, *** = p<0.001, **** = p<0.0001)

To further characterize the association of transcription activation to subnuclear structures during E_2_ stimulation, we employed our previously established *NRIP1* 24xMS2 RNA visualization cell line and virally transduced SC35 and Matrin3 fused to a florescent tag **(Fig. 5c)**. In line with DNA FISH data, RNA timelapse of *NRIP1* showed peak induced proximity to the periphery of interchromatin granules during acute E_2_ induction **(Fig. 5d).** This same phenomenon was observed for *TFF1* via immuno-RNA FISH **(Supplemental Fig. 3c)**.

We propose that induced association between activated ERα loci and subnuclear bodies may directly impact DNA mobility and target gene bursting. To examine this hypothesis, we performed three-color living imaging, combining RNA and DNA visualization platforms with an endogenous fluorescent tag of SC35 **(Fig. 5e; Supplemental Movie 6).** Surprisingly, we found no correlation between *NRIP1* DNA mobility and proximity to SC35, nor did we see any correlation between SC35 proximity and *NRIP1* gene bursting **(Fig. 5f).** Thus, while acutely activated ERα loci do change their sub-nuclear body associations with increased proximity in response to ligand stimulation, in line with previous finding, these associations appear to not have a direct impact upon either target gene bursting itself or DNA mobility of the regulated locus, suggesting functional importance on a higher order, macro level of architectural association.

### ERα and DNA kinetics during burst events as a surrogate for detecting condensate formation

To investigate whether the decreased DNA kinetics during bursting might reflect condensate formation in the signal-activated transcription unit, we elected to visualize ERα and DNA kinetics under conditions where bursting was removed from the equation by use of flavopiridol. We first confirmed that treatment with flavopiridol pauses transcription by >90% in our E_2_-induced WT MCF7 and *NRIP1* DNA cell lines **(Fig. 6a)**. In the absence of burst events, we hypothesized that DNA mobility would be increased, similar to the phenomenon seen in the chronic stimulated state, where burst frequency was half that of the acute state. In line with our prediction, visualization data revealed that *NRIP1* DNA motility was indeed faster with flavopiridol treatment **(Fig. 6b)**.

**Figure 6.**
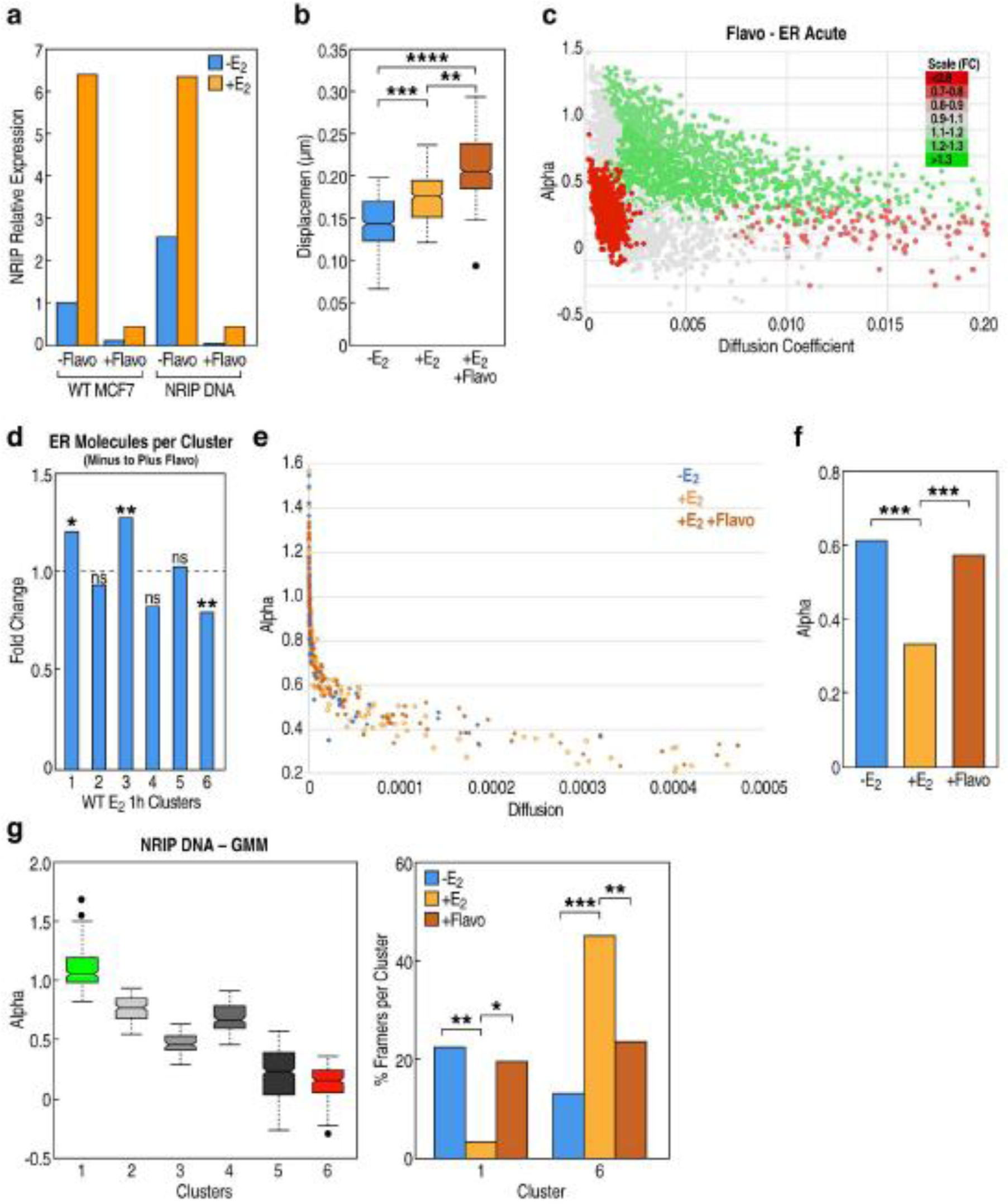
DNA kinetics as a surrogate for detecting condensate formation during burst events. **a.** Transcriptional pausing of *NRIP1* via flavopiridol treatment in WT MCF7 and *NRIP1* DNA visualization cell line measured by qPCR. **b.** Increased in DNA mobility post flavopiridol treatment detected at *NRIP1* loci. (n=68) **c.** SMT detects a significant depletion of previously identified acute specific, 1,6-hexanidiol sensitive ERα cluster post flavopiridol treatment. **d.** Bar graphs show fold change of ERα molecules per cluster post flavopiridol, with most confined cluster 6 showing the greatest depletion and less restricted clusters 1 and 3 gaining in ERα molecules. **e.** Alpha vs diffusion for ERα molecules in E_2_ minus, acute, and acute plus flavopiridol conditions. **f.** Flavopiridol treatment results in a statistical gain in alpha for ERα molecules. **g.** GMM clustering showed statistical significant gains and losses in the most confined (cluster 6) and least restricted (cluster 1) clusters in response to E_2_ stimulation and flavopiridol induced transcriptional pausing. (All statistics: 2-Tailed Paired T Test * = p<0.05, ** = p<0.01, *** = p<0.001, **** = p<0.0001)

We next wanted to determine whether there were changes to ERα kinetics in the absence of burst activation. To our surprise, we observed significant depletion of ERα molecules in the same 1,6-hexandiol sensitive fraction that we had identified **(Fig. 6c)**. The low diffusion, high confinement ERα population became less restricted as it gains in alpha and diffusion upon flavopiridol treatment, suggesting a similar disruption to condensate formation as had occurred with 1,6-hexandiol treatment **(Fig. 6d)**.

Given that alpha is a measure of confinement and a better proxy for condensate formation while diffusion coefficient is a speed parameter, we wanted to gain specific insight into alpha with our *NRIP1* DNA kinetics plus and minus flavopiridol. In support of a low sub-diffusive confined state during burst activation, we observed a decrease in alpha upon E_2_ stimulation. Consistently, upon transcriptional pausing with flavopiridol, there was a particular gain in alpha **(Fig. 6e, 6f)**. This data indicate that DNA kinetics become unrestricted in the absence of bursting, in support of condensate formation during gene activation events. Because the literature has reported heterogeneity of burst events across cells, we used GMM clustering on visualized DNA tracks to further validate the role of alpha and thus condensates during E_2_-induced transcription activation. Two clusters were of particular note, with a high fraction of unrestricted DNA tracks becoming heavily confined upon ligand stimulation, while flavopiridol loosened that restriction and shifted DNA tracks back into a non-confined state **(Fig. 6g)**. Therefore, differential DNA kinetics during burst vs non-burst vs no burst has emerged as a surrogate for detecting condensate formation during gene activation events, providing particular insight into regions of confinement on a micro level.

## Discussion

Our findings add important clarifications of the basis for increased mobility of regulated loci during gene activation. Here, we observe that activation, as defined by stimulation of a loci through either a developmental or environmental signal, not only results in increased displacement but consists of two distinct mobility phases: a high mobility, non-burst phase, and a lower-mobility burst phase. Previous studies were blind to bursting dynamics and therefore were constrained to identify a single high-mobility phase that was a mixture of the burst and non-burst phases^55^. We now can visualize that the highest-mobility state is constricted to the non-burst phase of a stimulated loci. While the full scope of the alterations in the enhancer condensates underlying and the functional consequences of this high mobility phase remain incompletely known, the increased mobility could conceivably facilitate increased frequency of interaction between activating elements such as enhancers and thus serve as a kinetic enhancer of stochastic transcriptional processes. Thus, against our initial expectations, active transcription is not required for the highest mobility state of the regulated genomic locus. Similar kinetic facilitation has been observed in B-cell recombination^56,57^. In contrast, we propose that the reduction in mobility observed during the burst-phase indicates that bursting places an activated loci into a fundamentally different kinetic state, governed by fundamentally different interactions between co-regulators and chromatin, and state of the MegaTrans condensate, and potentially interactions with subnuclear architectural machinery, presumably imparting a functional effect on enhancer:promoter interaction-mediated transcription.

In our exploration of what may be mechanistically responsible for the reduced mobility during bursting, we were surprised to see that association with both the nuclear matrix and the interchromatin granule seemed to have no kinetic impact upon *NRIP1* chromatin mobility, nor did association with the interchromatin granule significantly correlate with gene bursting. Thus, while the body of literature associating these organelles with gene transcription clearly supports their general functional importance in transcription REFS, our results suggest these organelles may play a more indirect role in the facilitation of bursting. The single molecule tracking (SMT) of glucocorticoid receptor (GR) has been recently used to show that acute versus chronic GR stimulation causes alterations in the bound fraction and binding duration of GR molecule, though the low temporal resolution of the SMT permits a somewhat limited insight into the detailed biophysical changes that occur to GR in chronic stimulation^40^. Furthermore, higher temporal resolution SMT of GR indicated the existence of a non-chromatin bound but low-mobility fraction of GR, with reliance of this fraction upon the GR intrinsically disordered domain^58^.

Conversely, we observe that the single molecule dynamics of ERα, which is both a MegaTrans-condensate component as well as the key transcription factor responsible for *NRIP1* activation, do in fact change in a distinct manner associated with <u>reduced</u> mobility in a state correlated with higher bursting rates. Given the observation in previous studies that ERα and other nuclear receptors bind to regulated gene loci and their enhancers, and that such nuclear receptor binding corresponds to bursts of gene transcription^25^, we speculate that the increased fraction of low-mobility ERα observed in the high-burst acute condition may facilitate both higher bursting rates and reduced chromatin mobility. Furthermore, our data showing the particular sensitivity of the low-mobility ERα fraction to 1,6-hexanediol suggests that this population of ERα molecules may exhibit condensates with potentially LLPS-like properties.

This model of a highly-concentrated, localized condensate of transcription factors facilitating transcription would fit current models of transcription being governed by the transient formation of co-factor-rich transcription hubs. In this model, enhancers and regulatory elements of varying degrees of genomic distribution come together in 4-dimensional space to form a critical concentration of valency that is sufficient to support bursts of gene transcription^18,49,59–66^. This model first provides a functional purpose to the increased “search-phase” mobility seen during the non-burst phase, as this increased chromatin kinetics will probabilistically increase the frequency at which a critical number of enhancers come into sufficient proximity so as to facilitate condensate formation and transcriptional bursting. This model of condensate-associated bursting both explains the viscosity-based reduction in mobility observed during gene bursting, and the ability of a single condensate to activate multiple genes simultaneously, and without direct interaction, two properties beyond the classic “enhancer-promoter looping” models of transcription^67–69^. Our paper provides striking evidence of a novel chromatin mobility state associated with gene bursting, which supports the model of condensate driven, enhancer-regulated gene expression. Our results indicate a multi-state model of DNA mobility during enhancer regulated transcription: 1) a low-mobility and transcriptionally inactive state; 2) a poised, high-mobility, pre- and post-burst state; and 3) an intermediate mobility state that occurs during bursting and is associated with a low-mobility, 1,6-hexanediol-sensitive condensate. Given the dynamic nature of condensate formation and its role in regulated transcription, it evokes the challenge of further delineating the differential components and kinetics constituting a condensate during burst and non-burst states.

## Acknowledgements

The authors acknowledge and thank Janet Hightower for assistance with figure preparation. The authors would like to acknowledge UCSD Neuroscience Microscopy Core managed by Core Director Dr. Binhai Zheng and Managing Director Jennifer Santini (NINDS P30NS047101). The authors would like to acknowledge UCSD Nikon Imaging Center previously managed by Director Dr. Eric Griffis and now under the management of Director Dr. Peng Guo. The authors would like to acknowledge UCSD Human Embryonic Stem Cell Core managed by Core Director Dr. Karl Willert and Manger Cody Fine. S.W was supported by a T32 training grant awarded to Dr. Brody (5T32AI007469-28). These studies were supported by grants from NIDDK (RO1DK018477, RO1DK039949) to M.G.R. M.G.R. was an investigator with the Howard Hughes Medical Institute in the course of these studies.

## Author Contributions

S.W., T.S., and M.G.R. designed the experimental strategies. S.W. and T.S. performed and analyzed the live and fixed cell imaging experiments. S.W. and T.S. performed the vector design, and together with Y.K. performed the cloning. A.G., S.J.N., and S.O. performed the PRO-seq assays and informatics analysis. A.G. and F.Y. performed the ChIP-seq assay and informatics analysis. K.O. prepared the samples for deep sequencing. T.W. assisted on the flavopiridol experiments and S.G. performed the qPCR experiments.

The authors declare no competing interests.

---

**Supplemental Figure 1.**
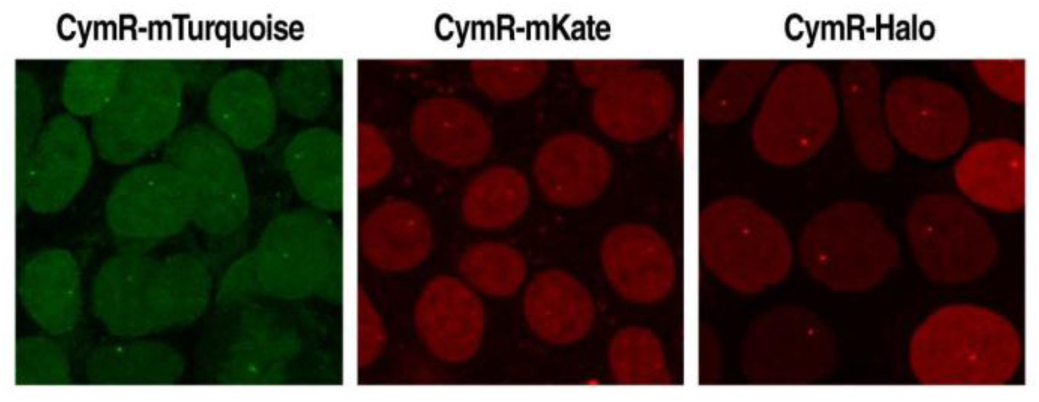
DNA visualization of *NRIP1* via CymR protein fused to either mTurquoise2, mKate, or Halo JaneliaFluor-549.

**Supplemental Figure 2.**
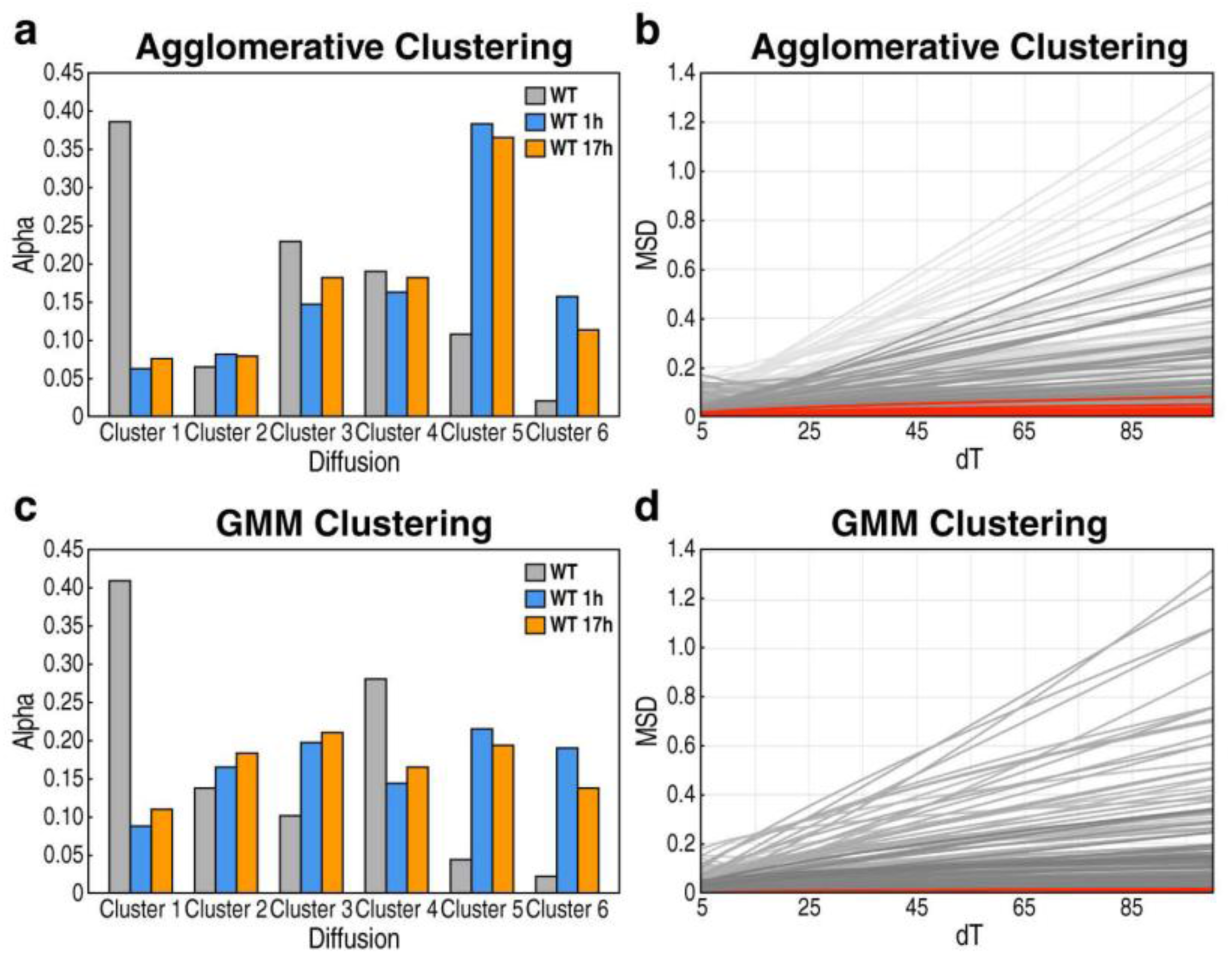
**ERα SMT**. **a.** Agglomerative clustering identified 6 distinct ERα populations characterized by alpha and diffusion in minus E2, acute, and chronic states. MSD curves of a randomized subset of single molecules from each cluster are graphed. **b.** GMM clustering similarly identified 6 distinct ERα populations characterized by alpha and diffusion in minus E2, acute, and chronic states. MSD curves of a randomized subset of single molecules from each cluster are graphed.

**Supplemental Figure 3.**
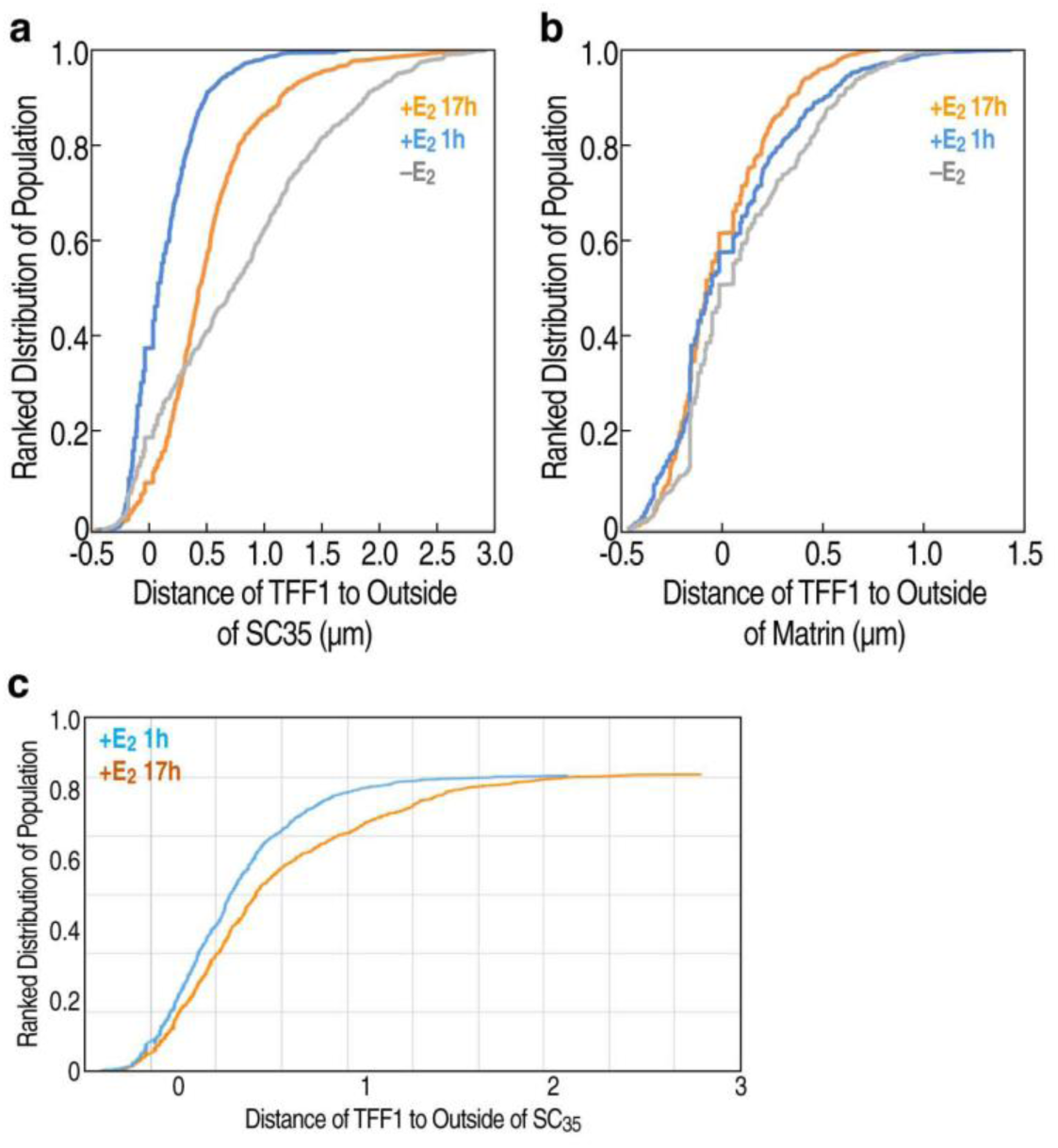
**TFF1 FISH**. **a.** Immuno-DNA FISH of *TFF1* shows increased proximity to SC35 in acute phase relative to chronic. **b.** Immuno-DNA FISH of *TFF1* shows induced association with Matrin3 network in the chronic state relative to acute. **c.** Immuno-RNA FISH of *TFF1* shows increased proximity to SC35 in acute relative to chronic phase.

**Supplemental Movie 1.** *NRIP1* RNA visualization timelapse of burst events.

**Supplemental Movie 2.** *NRIP1* DNA visualization timelapse of mobility.

**Supplemental Movie 3.** Single molecule timelapse of ERα in minus E_2_ condition.

**Supplemental Movie 4.** Single molecule timelapse of ERα in acute E_2_ condition.

**Supplemental Movie 5.** Single molecule timelapse of ERα in chronic E_2_ condition.

**Supplemental Movie 6.** Segmented timelapse of *NRIP1* RNA, DNA, and interchromatin granule 3-color visualization.

## Methods and Materials

### Antibodies

The antibodies in this study were:

anti-ER (Santa Cruz, HC-20), anti-GATA3 (Santa Cruz, HG3-31), Anti-RARa (Diagenode, C15310155), anti-AP2g (Santa Cruz, H-77), anti-SON (Abcam, ab121759), anti-Matrin3 (Abcam, ab70336).

### Tissue culture and treatment

Wild-type mammalian MCF7 breast cancer cells were cultured in High Glucose DMEM plus GlutaMAX (Gibco/Thermo Fisher Scientific) supplemented with 10% FBS (Omega) in a humidified incubator at 37^0^C with 5% CO2. Engineered stable cell line cultures were further supplemented with 1000x Gentamicin (50mg/mL, Omega) and 100x Amphotericin B (Gemini Bio).

To induce estrogen-mediated signaling, cells were washed and cultured in pheno-red free DMEM white medium (Gibco/Thermo Fisher Scientific) supplemented with 5% charcoal stripped FBS (Omega). The cells were then treated with 100nM 17β-estradiol (E_2_; Steraloids, Inc) for indicated time points. For 1,6-hexandiol treatment, cells were treated at 2%, 3%, 4%, and 5% (Sigma) in pheno-red free DMEM with 5% charcoal stripped FBS for 5minutes prior to washing 2x. The ethanol (EtOH) vehicle control was 0.05% in ChIP-seq and ICI 182780 (Abcam) control was used instead of EtOH for PRO-seq samples. Cells were treated drugs (EtOH or E_2_) for 1 hour or 16 hours for ChIP-seq or PRO-seq assays. For all experiments involving transcriptional pausing, cells where treated with 5uM of flavopiridol (Sigma) 1 hour prior to E_2_ stimulation.

### Plasmid generation

#### CRISPR cas9 plasmids

RNA_NRIP3’UTR_gRNA ATGTACCTGCCATCCAGTTTTGG was cloned into pSpCas9-PX459 V2.0 (Addgene 62988) using BbsI. The following primers were designed for gRNA hybridization:

RNA_NRIP3’U_gRNA_F
CACCG ATGTACCTGCCATCCAGTTT

RNA_NRIP3’U_gRNA_R
AAAC AAACTGGATGGCAGGTACAT C

DNA_NRIP_gRNA GCTATCAGTCGCTGCACCGAAGG was cloned into pSpCas9-PX459 V2.0 using BbsI. The following primers were designed for gRNA hybridization:

DNA_NRIP_gRNA_F
CACC GCTATCAGTCGCTGCACCGA

DNA_NRIP_gRNA_R
AAAC TCGGTGCAGCGACTGATAGC

SMT_ESR1_gRNA GGGAGCTCTCAGACCGTGGCAGG was cloned into pSpCas9-PX459 V2.0 using BbsI. The following primers were designed for gRNA hybridization:

ER_gRNA_F
CACC GGGAGCTCTCAGACCGTGGC

ER_gRNA_R
AAAC GCCACGGTCTGAGAGCTCCC

SRSF2_gRNA TAGCTCATAGCTCTGAGTGGCGG was cloned into pSpCas9-PX459 V2.0 using BbsI. The following primers were designed for gRNA hybridization:

SC35_gRNA_F
CACCG TAGCTCATAGCTCTGAGTGG

SC35_gRNA_R
AAAC CCACTCAGAGCTATGAGCTA C

#### RNA visualization donor plasmid

CRISPR cas9 was used knock-in 24xMS2 hairpins into 3’UTR of *NRIP1.* To generate NRIP1-24xMS2 RNA donor vector, cloning was completed in 4 steps: 1.) PCR left *NRIP1* homology arm, 2.) PCR right *NRIP1* homology arm, 3.) PCR HaloTag, 4.) Gibson assembly left and right homology arms with HaloTag, and 5.) insert 24xMS2 hairpins at corresponding digestion sites. The left and right homology arms of *NRIP1* was designed around NRIP1-gRNA sequence, and has that same NRIP-gRNA flanking both sides of the arms to induce simultaneous double stranded break of donor plasmid as well as *NRIP1* locus to efficiently induce HDR of linearized 24xMS2 hairpins. The homology arms were PCR amplified using AccuPrime Pfx SuperMix (Thermo Fisher) standard protocol from MCF7 genomic DNA and cloned into pmCherry-N1 (Addgene Plasmid #54517) to flank HaloTag and 24xMS2 hairpins. The left arm, right arm, and HaloTag were assembled via HiFi Gibson Assembly (NEB E2621L) following recommended protocol from NEB for multi-fragment assembly. mCherry-C1 plasmid backbone was digested with EcoRI (NEB) and BamHI (NEB) prior to Gibson assembly, and destroyed upon successful assembly. mCherry tag is used during the first round of flow cytometry to enrich for MCF7 cells that have been successfully transfected donor plasmid. The HaloTag knocked-in right in front of the stop codon for *NRIP1* was used to enriched for 24xMS2 stable cell lines in the second round of flow cytometry. The following primers and templates were used for left and homology arms as well as HaloTag with appropriate overhangs for Gibson in later steps (corresponding sequences for PCR amplification in bold, gRNA in italics):

NRIP_LeftArm_F
CCGGCGGCATGGACGAGCTGTACAAGGAATAG*ATGTACCTGCCATCCAGTTTTGG* **CAACATTGTTGATGCTGCAAACAATCACAGTGCCC**

NRIP_LeftArm_R
TTCTGATTCTTTCTTTATCGTTAGCACGCTTCCCAGAAGTCC

NRIP_LeftArm
**CAACATTGTTGATGCTGCAAACAATCACAGTGCCC**CAGAAGTACTGTATGGGTCC TTGCTTAACCAGGAAGAGCTGAAATTTAGCAGAAATGATCTTGAATTTAAATATCCTGCTGGTCATGGCTCAGCCAGCGAAAGTGAACACAGGAGTTGGGCCAGAGAGAGCAAAAGCTTTAATGTTCTGAAACAGCTGCTTCTCTCAGAAAACTGTGTGCGAGATTTGTCCCCGCACAGAAGTAACTCTGTGGCTGACAGTAAAAAGAAAGGACACAAAAATAATGTGACCAACAGCAAACCTGAATTTAGCATTTCTTCTTTAAATGGACTGATGTACAGTTCCACTCAGCCCAGCAGTTGCATGGATAACAGGACATTTTCATACCCAGGTGTAGTAAAAACTCCTGTGAGTCCTACTTTCCCTGAGCACTTGGGCTGTGCAGGGTCTAGACCAGAATCTGGGCTTTTGAATGGGTGTTCCATGCCCAGTGAGAAAGGACCCATTAAGTGGGTTATCACTGATGCGGAGAAGAATGAGTATGAAAAAGACTCTCCAAGATTGACCAAAACCAACCCAATACTATATTACATGCTTCAAAAAGGAGGCAATTCTGTTACCAGTCGAGAAACACAAGACAAGGACATTTGGAGGGAGGCTTCATCTGCTGAAAGTGTCTCACAGGTCACAGCCAAAGAAGAGTTACTTCCTACTGCAGAAACGAAAGCTTCTTTCTTTAATTTAAGAAGCCCTTACAATAGCCATATGGGAAATAATGCTTCTCGCCCACACAGCGC AAATGGAGAAGTTTAT**GGACTTCTGGGAAGCGTGCTAACGATAAAGAAAGAATC AGAA**

NRIP_RightArm_F
**TTTTGGATCTTTTTAAAACTAATGAGTATGAACTTGAGATCTGTATAAATAAGA GCATG**

NRIP_RightArm_R
GCTGATTATGATCAGTTATCTAGATCCGGTG*ATGTACCTGCCATCCAGTTTTGG* **CATAACACTCGCAGGTACCACAGGAATGGTGAG**

NRIP_RightArm
**TTTTGGATCTTTTTAAAACTAATGAGTATGAACTTGAGATCTGTATAAATAAGA GCATG**ATTTGAAAAAAAGCATGGTATAATTGAAACTTTTTTCATTTTGAAAAGTATT GGTTACTGGTGATGTTGAAATATGCATACTAATTTTTGCTTAACATTAGATGTCATGAGGAAACTACTGAACTAGCAATTGGTTGTTTAACACTTCTGTATGCATCAGATAACAACTGTGAGTAGCCTATGAATGAAATTCTTTTATAAATATTAGGCATAAATTAAAATGT AAAACTCCATTCATAGTGGATTAATGCATTTTGCTGCCTTTATTAGGGTACTTTATTTTGCTTTTCAGAAGTCAGCCTACATAACACATTTTTAAAGTCTAAACTGTTAAACAACTCTTTAAAGGATAATTATCCAATAAAAAAAAACCTAGTGCTGATTCACAGCTTATTATCCAATTCAAAAATAAATTAGAAAAATATATGCTTACATTTTTCACTTTTGCTAAAAAGAAAAAAAAAAGGTGTTTATTTTTAACTCTTGGAAGAGGTTTTGTGGTTCCCAATGTGTCTGTCCCACCCTGATCCTTTTCAATATATATTTCTTTAAACCTTGTGCTACTTAGTAAAAATTGATTACAATTGAGGGAAGTTTGATAGATCCTTTAAAAAAAAGGCAGATTTCCATTTTTTGTATTTTAACTACTTTACTAAATTAATACTCCTCCTTTTACAGAATTAGAAAAGTTAACATTTATCTTTAGGTGGTTTCCTGAAAAGTTGAATATTTAAGAAATTGTTTTTAACAGAAGCAAAATGGCTTTTCTTTGGACAGTTTTCACCATCTCTTGTAAAAG TTAATT**CTCACCATTCCTGTGGTACCTGCGAGTGTTATG**

NRIP_Halo_F
GAAGCGTGCTAACGATAAAGAAAGAATCAGAA
**ATGGCAGAAATCGGTACTGGCTTTCCATTCGACCCC**

NRIP_Halo_R
CATACTCATTAGTTTTAAAAAGATCCAAAAGGATCC CTGGATGGCAGGTACATTTTA
**GCCGGAAATCTCGAGCGTCGACAGCCAG**

pENTR4-HaloTag (Addgene 29644)
**ATGGCAGAAATCGGTACTGGCTTTCCATTCGACCCC**CATTATGTGGAAGTCCTGG GCGAGCGCATGCACTACGTCGATGTTGGTCCGCGCGATGGCACCCCTGTGCTGTTCCTGCACGGTAACCCGACCTCCTCCTACGTGTGGCGCAACATCATCCCGCATGTTGCACCGACCCATCGCTGCATTGCTCCAGACCTGATCGGTATGGGCAAATCCGACAAACCAGACCTGGGTTATTTCTTCGACGACCACGTCCGCTTCATGGATGCCTTCATCGAAGCCCTGGGTCTGGAAGAGGTCGTCCTGGTCATTCACGACTGGGGCTCCGCTCTGGGTTTCCACTGGGCCAAGCGCAATCCAGAGCGCGTCAAAGGTATTGCATTTATGGAGTTCATCCGCCCTATCCCGACCTGGGACGAATGGCCAGAATTTGCCCGCGAGACCTTCCAGGCCTTCCGCACCACCGACGTCGGCCGCAAGCTGATCATCGATCAGAACGTTTTTATCGAGGGTACGCTGCCGATGGGTGTCGTCCGCCCGCTGACTGAAGTCGAGATGGACCATTAC CGCGAGCCGTTCCTGAATCCTGTTGACCGCGAGCCACTGTGGCGCTTCCCAAACGAGCTGCCAATCGCCGGTGAGCCAGCGAACATCGTCGCGCTGGTCGAAGAATACATGGACTGGCTGCACCAGTCCCCTGTCCCGAAGCTGCTGTTCTGGGGCACCCCAGGCGTTCTGATCCCACCGGCCGAAGCCGCTCGCCTGGCCAAAAGCCTGCCTAACTGCAAGGCTGTGGACATCGGCCCGGGTCTGAATCTGCTGCAAGAAGACAACCCGGACCTGATCGGC AGCGAGATCGCGCG**CTGGCTGTCGACGCTCGAGATTTCCGGC**

Upon Sanger sequencing to validate successful insertion of homology arms with HaloTag in mCherry-C1 plasmid backbone, BamHI was used to digest newly sequenced backbone, and BamHI and BlgII (NEB) was used to extract 24xMS2 from pET259-pUC57-24xMS2V6 (Addgene 104391). Sanger sequencing was completed again to determine the correct orientation of 24xMS2 hairpins in donor plasmid.

24xMS2V6
GGATCCCAGAGCCCCCTGGCAATCGCGGGGAGACGCAGGACTACCGCGTCTTCACTCTCGCTTGCGCGGTATACGCAGGGAGCAACGTCACCCAGGGCGACACAGAGGAATACCCTGTGTGGCCCCACGGTGCTACAAAGAACATTCCGTATTGCTCACCAGTGTCACATGTGGAGGACTACCCCACAAGCGAGTCAGGAAACCTTCGGGGCATCGCACCATTATCCCGAACAATCGACACGCAGGATTACCGCGTGGGACACTCTGTTCCCCGTCAAAATTGGACCATACCGGAGTCGGCTTACGTCATGCAGGATTACCGCATGCATGGTGCAGAATATCGGCATGTCTACTCGTACAGTCAAATCTACTCGTGTGGTCAGGACTACCGACCACTTCTATTCTATTCATCTTTTCGGTTGTGCAGGTATTCTGCCGATGTACGAGAAGACGATTACGCTTCTCGACTACCTTCATCATACATGGTGTGCAGATGGCCGCCAAGTTTTTTGCCAATGGAGGAATACCCCATTCTCTGTCAACCAACCCATGCAAAGTTTACACTCTGCTATGGCAGCACTGTCGCAGAGGAATACCCTGCGAGCCAAAACGGCCCCCGGTGCGTGTATTGCATCTGCCTTGCGAGCATTCACAGGGACGAATACGCCCTGCCGTTGCATTACTTCAATATGGGTGCTCTGTCGTCGTCATCAGGACCATTTGCGCAGGACTACCGCGCATATATCATCAGCACTCGTGCGGATACTTCTGGGATTCCTATTGTTACGCGAGCTCAGGAATACCGAGCTCTGGCGACAGAGACCCTCACACGGAAGATCCCAGAGCCCCCTGGCAATCGCGGGGAGACGCAGGACTACCGCGTCTTCACTCTCGCTTGCGCGGTATACGCAGGGAGCAACGTCACCCAGGGCGACACAGAGGAATACCCTGTGTGGCCCCACGG TGCTACAAAGAACATTCCGTATTGCTCACCAGTGTCACATGTGGAGGACTACCCCACAAGCGAGTCAGGAAACCTTCGGGGCATCGCACCATTATCCCGAACAATCGACACGCAGGATTACCGCGTGGGACACTCTGTTCCCCGTCAAAATTGGACCATACCGGAGTCGGCTTACGTCATGCAGGATTACCGCATGCATGGTGCAGAATATCGGCATGTCTACTCGTACAGTCAAATCTACTCGTGTGGTCAGGACTACCGACCACTTCTATTCTATTCATCTTTTCGGTTGTGCAGGTATTCTGCCGATGTACGAGAAGACGATTACGCTTCTCGACTACCTTCATCATACATGGTGTGCAGATGGCCGCCAAGTTTTTTGCCAATGGAGGAATACCCCATTCTCTGTCAACCAACCCATGCAAAGTTTACACTCTGCTATGGCAGCACTGTCGCAGAGGAATACCCTGCGAGCCAAAACGGCCCCCGGTGCGTGTATTGCATCTGCCTTGCGAGCATTCACAGGGACGAATACGCCCTGCCGTTGCATTACTTCAATATGGGTGCTCTGTCGTCGTCATCAGGACCATTTGCGCAGGACTACCGCGCATATATCATCAGCACTCGTGCGGATACTTCTGGGATTCCTATTGTTACGCGAGCTCAGGAATACCGAGCTCTGGCGACAGAGACCCTCACACGGAAGATCT

#### DNA visualization donor plasmid

DNA donor plasmid for integrase knock-in was designed according to protocol described in Alexander et al. 2019. The following duplexed ultramer was designed to first knock-in Bcb1 integrase landing pad adjacent to *NRIP* locus centered around gRNA chosen for DNA visualization described above (landing pad in bold, gRNA in italics):

NRIP1_Bxb1
CTATATTCTTTCTAGGTGTTAGGAATGCTCCCAGGAGGTACAAAA*GCTATCAGTCGCT GCACCGA*
**tctagagtcgtggtttgtctggtcaaccaccgcggtctcagtggtgtacggtacaaaccccgacctcgag***AGG*ACTCTAGG ATGCTATGACTTAGTGATTATCTCTGCCTTTTGTAATTTCTCTCTCTCCCCCTT

Following successful knock-in of landing pad via CRISPR, Bxb1 integrase (Addgene 119901) will cut and knock-in 144xCuO (Addgene 119903) array at site of landing pad.

#### RNA + DNA visualization donor plasmid

RNA donor plasmid for RNA+DNA dual visualization was slightly modified from method described above under “RNA visualization donor plasmid”. mCherry was first removed from pC1 donor plasmid. To remove sequence, mCherry-pC1 was digested using AgeI (NEB) and XhoI (NEB). The following primers were designed for hybridization, following by ligation to pC1 digested backbone:

pC1_F
CCGGT AGCCATGGCTGTGCAGCTT C

pC1_R
TCGAG AAGCTGCACAGCCATGGCT A

Previous steps to create left and right *NRIP1* homology arms were repeated to recreate 24xMS2 *NRIP1* donor plasmid in newly created blank pC1 backbone. Rather than HaloTag, however, puromycin tag was used instead. The following primers were used to amplify puromycin, and includes the appropriate linker and overhangs for Gibson assembly (corresponding sequences for PCR amplification in bold):

NRIP1_puro_F
GAAGCGTGCTAACGATAAAGAAAGAATCAGAAggtggttctggtggtggttctggt
**ATGACCGAGTACAAGCCCACGGTGCGCC**

NRIP1_puro_R
CATACTCATTAGTTTTAAAAAGATCCAAAAGGATCC CTGGATGGCAGGTACATTTTA
**ACCGGGCTTGCGGGTCATGCACCAGG**

Puromycin (Addgene 62988)
**ATGACCGAGTACAAGCCCACGGTGCGCC**TCGCCACCCGCGACGACGTCCCCCGG GCCGTACGCACCCTCGCCGCCGCGTTCGCCGACTACCCCGCCACGCGCCACACCGTCGACCCGGACCGCCACATCGAGCGGGTCACCGAGCTGCAAGAACTCTTCCTCACGCGCGTCGGGCTCGACATCGGCAAGGTGTGGGTCGCGGACGACGGCGCCGCGGTGGCGG TCTGGACCACGCCGGAGAGCGTCGAAGCGGGGGCGGTGTTCGCCGAGATCGGCCCGCGCATGGCCGAGTTGAGCGGTTCCCGGCTGGCCGCGCAGCAACAGATGGAAGGCCTCCTGGCGCCGCACCGGCCCAAGGAGCCCGCGTGGTTCCTGGCCACCGTCGGCGTCTCGCCCGACCACCAGGGCAAGGGTCTGGGCAGCGCCGTCGTGCTCCCCGGAGTGGAGGCGGCCGAGCGCGCCGGGGTGCCCGCCTTCCTGGAGACCTCCGCGCCCCGCAACCTCCCCTTCTACGAGCGGCTCGGCTTCACCGTCACCGCCGACGTCGAGGTGCCCGAAGGAC CGCGCA**CCTGGTGCATGACCCGCAAGCCCGGT**

#### ESR1 SMR visualization donor plasmid

ERα donor plasmid was cloned using the same methods as described above under “RNA visualization donor plasmid”. For SMT, HaloTag was knocked into the ESR1 locus directly in front of the stop codon. The following primers were used for right and left ESR1 homology arms and HaloTag with appropriate overhangs for Gibson Assembly (corresponding sequences for PCR amplification in bold, gRNA in italics):

ESR1_LeftArm_F
GGGCCGCCACTCCACCGGCGGCATGGACGAGCTGTACAAGGAATTCTAG*GGGAGCT CTCAGACCGTGGCAGG* **CCCCATACTCTATTCCGAGTATGATCCTACCAGACCC**

ESR1_LeftArm_R
AATGGGGGTCGAATGGAAAGCCAGTACCGATTTCTGCCAT **GACCGTGGCAGGGAAACCCTCTGCCTCC**

ESR1_LeftArm
**CCCCATACTCTATTCCGAGTATGATCCTACCAGACCC**TTCAGTGAAGCTTCGATG ATGGGCTTACTGACCAACCTGGCAGACAGGGAGCTGGTTCACATGATCAACTGGGCGAAGAGGGTGCCAGGCTTTGTGGATTTGACCCTCCATGATCAGGTCCACCTTCTAGAATGTGCCTGGCTAGAGATCCTGATGATTGGTCTCGTCTGGCGCTCCATGGAGCACCCAGGGAAGCTACTGTTTGCTCCTAACTTGCTCTTGGACAGGAACCAGGGAAAATGTGTAGAGGGCATGGTGGAGATCTTCGACATGCTGCTGGCTACATCATCTCGGTTCCGCATGATGAATCTGCAGGGAGAGGAGTTTGTGTGCCTCAAATCTATTATTTTGCTTAATTCT GGAGTGTACACATTTCTGTCCAGCACCCTGAAGTCTCTGGAAGAGAAGGACCATATCCACCGAGTCCTGGACAAGATCACAGACACTTTGATCCACCTGATGGCCAAGGCAGGCCTGACCCTGCAGCAGCAGCACCAGCGGCTGGCCCAGCTCCTCCTCATCCTCTCCCACATCAGGCACATGAGTAACAAAGGCATGGAGCATCTGTACAGCATGAAGTGCAAGAACGTGGTGCCCCTCTATGACCTGCTGCTGGAGATGCTGGACGCCCACCGCCTACATGCGCCCACTAGCCGTGGAGGGGCATCCGTGGAGGAGACGGACCAAAGCCACTTGGCC ACTGCGGGCTCTACTTCATCGCATTCCTTGCAAAAGTATTACATCACGGG**GGAGGC AGAGGGTTTCCCTGCCACGGTC**

ESR1_RightArm_F
CGAGATCGCGCGCTGGCTGTCGACGCTCGAGATTTCCGGC **TGAGAGCTCCCTGGCTCCCACACGGTTCAG**

ESR1_RightArm_R
ATGTGGTATGGCTGATTATGATCAGTTATCTAGATCCGGTGGATCC *GGGAGCTCTCAGACCGTGGCAGG* **CAGGAACTTATCCCTCATATAGGGAGACTTAACTAAGATCAACTTGCTG**

ESRA_RightArm
**TGAGAGCTCCCTGGCTCCCACACGGTTCAG**ATAATCCCTGCTGCATTTTACCCTC ATCATGCACCACTTTAGCCAAATTCTGTCTCCTGCATACACTCCGGCATGCATCCAACACCAATGGCTTTCTAGATGAGTGGCCATTCATTTGCTTGCTCAGTTCTTAGTGGCACATCTTCTGTCTTCTGTTGGGAACAGCCAAAGGGATTCCAAGGCTAAATCTTTGTAACAGCTCTCTTTCCCCCTTGCTATGTTACTAAGCGTGAGGATTCCCGTAGCTCTTCACAGCTGAACTCAGTCTATGGGTTGGGGCTCAGATAACTCTGTGCATTTAAGCTACTTGTAGAGACCCAGGCCTGGAGAGTAGACATTTTGCCTCTGATAAGCACTTTTTAAATGGCTCTAAGAATAAGCCACAGCAAAGAATTTAAAGTGGCTCCTTTAATTGGTGACTTGGAGAAAGCTAGGTCAAGGGTTTATTATAGCACCCTCTTGTATTCCTATGGCAATGCATCCTTTTATGAAAGTGGTACACCTTAAAGCTTTTATATGACTGTAGCAGAGTATCTGGTGATTGTCAATTCATTCCCCCTATAGGAATACAAGGGGCACACAGGGAAGGCAGATCCCCTAGTTGGCAAGACTATTTTAACTTGATACACTGCAGATTCAGATGTGCTGAAAGC
TCTGCCTCTGGCTTTCCGGTCATGGGTTCCAGTTAATTCATGCCTCCCATGGACCTAT GGAGAG**CAGCAAGTTGATCTTAGTTAAGTCTCCCTATATGAGGGATAAGTTCCT G**

ESR1_Halo_F
TTACATCACGGGGGAGGCAGAGGGTTTCCCTGCCACGGTC **ATGGCAGAAATCGGTACTGGCTTTCCATTCGACCCC**

ESR1_Halo_R
CAGGGATTATCTGAACCGTGTGGGAGCCAGGGAGCTCTCA **GCCGGAAATCTCGAGCGTCGACAGCCAGC**

#### Interchromatin granule/SC35 donor plasmid

SC35 donor plasmid was cloned using similar technique as “RNA visualization donor plasmid” described above, but with restriction digest rather than Gibson Assembly into mCherry-pC1. mTurquoise2 was knocked into the SRSF2 locus directly in front of the stop codon. The left arm used EcoRI and KpnI (NEB). The right arm used KpnI and BamHI. mTurquoise2 was PCR amplified using AccuPrime Pfx with KpnI flanking both sides. Sanger sequencing was done to validate correct orientation of mTurquoise2 insertion. The following primers were used for right and left SRSF2 homology arms and mTurquoise2 with appropriate digestion sites (corresponding sequences for PCR amplification in bold, gRNA in italics):

SC35_LeftArm_F
ATATGAATTCTAG*TAGCTCATAGCTCTGAGTGGCGG* **GGAGCTCTCGGGCTCTGGCGAAAG**

SC35_LeftArm_R
ATTATGGTACC **AGCTCTGAGTGGCGGCCCGGAGC**

SC35_LeftArm
**GGAGCTCTCGGGCTCTGGCGAAAG**GGGGTTGGGAAGCGGACTCCGGCGAAGAGC AGTCAGGAACGGCTGGACGGGGCCGAGAGACGACGCAGCGGAGTCTGAGGGGGCCGGGGTCACAGGGAGAGGCAAATGAGGGCAGAAGCACTCTTCAGACGGAGACGGGGCTGACGGCTGAGCAAAAAAGGGGCGTGAACTTGGAGTATTGGCATGAAAAAAAGGTGGGGGCCCAGATGGGGAGCACCTCCTCTTCCTCCTGCCCAATCGCGATCGTCCGGCCTCCCAGGCGGAAAAAGCGTTTCGCGGGCTTTCCAACTGCCCGCTAATTCCGCTCCCCCTCCCTCCAGCCGTGACTCCGCGCTTTTTGGCCCGCCCGCCGGGCTGTGCGCAGGCGCTTCGGGTAGGGGCGGGGCGCGCGGCAGGGTCGTTACGAAGCGGGGCGCGGTGGGCCAATCAGAAGGTTTCATTTCCGGGTGGCGCGGGCGCCATTTTGTGAGGAGCGATATAAACGGGCGCAGAGGCCGGCTGCCCGCCCAGTTGTTACTCAGGTGCGCTAGCCTGCGGAGCCCGTCCGTGCTGTTCTGCGGCAAGGCCTTTCCCAGTGTCCCCACGCGGAAGGCAACTGCCTGAGAGGCGCGGCGTCGCACCGCCCAGAGCTGAGGAAGCCGGCGCCAGT TCGCGGG**GCTCCGGGCCGCCACTCAGAGCT**

SC35_RightArm_F
ATTATGGTACC **AGCTACGGCCGCCCCCCTCC**

SC35_RightArm_R
ATATGGATCC*TAGCTCATAGCTCTGAGTGGCGG* **GGTCGCAGACGGCGGAAGCTCGC**

SC35_RightArm
**AGCTACGGCCGCCCCCCTCC**CGATGTGGAGGGTATGACCTCCCTCAAGGTGGACA ACCTGACCTACCGCACCTCGCCCGACACGCTGAGGCGCGTCTTCGAGAAGTACGGGCGCGTCGGCGACGTGTACATCCCGCGGGACCGCTACACCAAGGAGTCCCGCGGCTTCGCCTTCGTTCGCTTTCACGACAAGCGCGACGCTGAGGACGCTATGGATGCCATGGACGGGGCCGTGCTGGACGGCCGCGAGCTGCGGGTGCAAATGGCGCGCTACGGCCGCCCCCCGGACTCACACCACAGCCGCCGGGGACCGCCACCCCGCAGGTACGGGGGCGGTGGCTACGGACGCCGGAGCCGCAGGTAAACGGGGCTGAGGGGACCGCGGGAGGCGGGGCGGGGCGCGCGGGAGGCCCGGGCGACCTCACAAAGGTCCGCGGCGAAGCACGTGGTGCGGGCCCGGACGGGGCGGGGGTGCACGCCGCGTCTCGCGACCCTCCGGCCAC CCC**GCGAGCTTCCGCCGTCTGCGACC**

mTurq_F
ATTATGGTACC **ATGGTGAGCAAGGGCGAGGAGCTG**

mTurq_R
ATTATGGTACC **CTTGTACAGCTCGTCCATGCCGAGAGTG**

mTurquoise2 (Addgene 54843) **ATGGTGAGCAAGGGCGAGGAGCTG**TTCACCGGGGTGGTGCCCATCCTGGTCGAGC TGGACGGCGACGTAAACGGCCACAAGTTCAGCGTGTCCGGCGAGGGCGAGGGCGATGCCACCTACGGCAAGCTGACCCTGAAGTTCATCTGCACCACCGGCAAGCTGCCCGTGCCCTGGCCCACCCTCGTGACCACCCTGTCCTGGGGCGTGCAGTGCTTCGCCCGCTACCCCGACCACATGAAGCAGCACGACTTCTTCAAGTCCGCCATGCCCGAAGGCTACGTCCAGGAGCGCACCATCTTCTTCAAGGACGACGGCAACTACAAGACCCGCGCCGAGGTGAAGTTCGAGGGCGACACCCTGGTGAACCGCATCGAGCTGAAGGGCATCGACTTCAAGGAGGACGGCAACATCCTGGGGCACAAGCTGGAGTACAACTACTTTAGCGACAACGTCTATATCACCGCCGACAAGCAGAAGAACGGCATCAAGGCCAACTTCAAGATCCGCCACAACATCGAGGACGGCGGCGTGCAGCTCGCCGACCACTACCAGCAGAACACCCCCATCGGCGACGGCCCCGTGCTGCTGCCCGACAACCACTACCTGAGCACCCAGTCCAAGCTGAGCAAAGACCCCAACGAGAAGCGCGATCACATGGTCCTGCTGGAGTTCGTG ACCGCCGCCGGGAT**CACTCTCGGCATGGACGAGCTGTACAAG**

#### Subnuclear structures viral constructs

All lentiviral vectors were created using pCDH-CMV-MCS-EF1α-Puro backbone (System Biosciences CD510B-1). To maintain a lower, uniform expression level for virally transduced subnuclear structures, CMV promoter was swapped out for PGK promoter from pCDH-PGK vector (Addgene 72268) using HpaI (NEB) and NheI (NEB) digestion followed by ligation.

To create pCDH-PGK-mTurq-SC35, mTurquoise2 fused to SRSF2 open reading frame with linker were first cloned with blank pC1 backbone via restriction digest followed by ligation to create pC1-mTurq-SC35. mTurquoise2 used EcoRI and XhoI while SRSF2 used XhoI and BamHI. The following primers were used (corresponding sequences for PCR amplification in bold):

mTurq_pC1_F
ATTATACCGGT **ATGGTGAGCAAGGGCGAGGAGCTG**

mTurq_pC1_R
ATTATCTCGAG **CTTGTACAGCTCGTCCATGCCGAGAGTG**

SC35_pC1_F
ATTATCTCGAG ggtggttctggtggtggttctggt **ATGAGCTACGGCCGCCCCCCTC**

SC35_pC1_R
ATTATGGATCC **TTAAGAGGACACCGCTCCTTCCTCTTC**

#### ORF: >gi|306482645:252-917 Homo sapiens serine/arginine-rich splicing factor 2 (SRSF2), transcript variant 2, mRNA

**ATGAGCTACGGCCGCCCCCCTC**CCGATGTGGAGGGTATGACCTCCCTCAAGGTGG ACAACCTGACCTACCGCACCTCGCCCGACACGCTGAGGCGCGTCTTCGAGAAGTACGGGCGCGTCGGCGACGTGTACATCCCGCGGGACCGCTACACCAAGGAGTCCCGCGGCTTCGCCTTCGTTCGCTTTCACGACAAGCGCGACGCTGAGGACGCTATGGATGCCATGGACGGGGCCGTGCTGGACGGCCGCGAGCTGCGGGTGCAAATGGCGCGCTACGGCCGCCCCCCGGACTCACACCACAGCCGCCGGGGACCGCCACCCCGCAGGTACGGGGGCGGTGGCTACGGACGCCGGAGCCGCAGCCCTAGGCGGCGTCGCCGCAGCCGATCCCGGAGTCGGAGCCGTTCCAGGTCTCGCAGCCGATCTCGCTACAGCCGCTCGAAGTCTCGGTCCCGCACTCGTTCTCGATCTCGGTCGACCTCCAAGTCCAGATCCGCACGAAGGTCCAAGTCCAAGTCCTCGTCGGTCTCCAGATCTCGTTCGCGGTCCAGGTCCCGGTCTCGGTCCAGGAGTCCTCCCCCAGTGTCCAAGAGGGAATCCAAATCCAGGTCGCGATCGA AGAGTCCCCCCAAGTCTCCT**GAAGAGGAAGGAGCGGTGTCCTCTTAA**

After validation of pC1-mTurq-SC35, NheI and BamHI was used to extract mTurq-SC35 from pC1 donor plasmid and transferred to pCDH-PGK backbone to create final lentiviral contruct pCDH-PGK-mTurq-SC35.

pCDH-PGK-mCardinal-Matrin was created using the same protocol as described for SC35, but rather than mTurquoise2 mCardinal was used instead. The following primers were used for mCardinal (corresponding sequences for PCR amplification in bold):

mCard_pC1_F
ATTATACCGGT **ATGGTGAGCAAGGGCGAGGAGCTGATC**

mCard_pC1_R
ATTATCTCGAG **CTTGTACAGCTCGTCCATGCCATTAAGTTTGTGC**

mCardinal (Addgene 54799)
**ATGGTGAGCAAGGGCGAGGAGCTGATC**AAGGAGAACATGCACATGAAGCTGTAC ATGGAAGGCACCGTGAACAACCACCACTTCAAGTGCACCACCGAAGGGGAGGGCAAGCCCTACGAGGGCACCCAGACCCAGAGGATTAAGGTGGTGGAGGGAGGCCCCCTGCCGTTCGCATTCGACATCCTGGCCACCTGCTTTATGTACGGGAGCAAGACCTTCATCAACCACACCCAGGGCATCCCCGATTTCTTTAAGCAGTCCTTCCCTGAGGGCTTCACATGGGAGAGAGTCACCACATACGAAGACGGGGGCGTGCTTACCGTTACCCAGGACACCAGCCTCCAGGACGGCTGCTTGATCTACAACGTCAAGCTCAGAGGGGTGAACTTCCCATCCAACGGCCCTGTGATGCAGAAGAAAACACTCGGCTGGGAGGCCACCACCGAGACCCTGTACCCCGCTGACGGCGGCCTGGAAGGCAGATGCGACATGGCCCTGAAGCTCGTGGGCGGGGGCCACCTGCACTGCAACCTGAAGACCACATACAGATCCAAGAAACCCGCTAAGAACCTCAAGATGCCCGGCGTCTACTTTGTGGACCGCAGACTGGAAAGAATCAAGGAGGCCGACAATGAGACCTACGTCGAGCAGCACGAGGTGGCTGTGGCCAGA TACTGCGACCTCCCTAGCAAACTGGG**GCACAAACTTAATGGCATGGACGAGCTGT ACAAG**

### Stable cell line generation

#### Endogenous RNA visualization

WT MCF7 were plated in 24 well at 75% confluency 1 day prior to transfection with 2ul Lipofectamine 2000 (Thermo Fisher) plus 0.3ug RNA_NRIP3’UTR_gRNA and 0.3ug of NRIP1-24xMS2 donor plasmid suspended in total of 100ul OptiMEM (Thermo Fisher). Cells were incubated in transfection reaction overnight and then recovered for 48 hours. On day 3 post transfection, transfected MCF7s were flow sorted for mCherry positive signal to enrich for cells that have taken up donor plasmid into 24 well. After 2 weeks of recovery, pooled population was resorted into single clones in 96 wells using Janelia Fluor HaloTag Ligand 646 (Promega GA1120) to enrich for clones with stable integration. PCR followed by Sanger sequencing was used to identify positive 24xMS2 hairpin insertion after 3 weeks of recovery. Upon validation of positive single clone, cell was then transfected again with MS2 coat protein MCP-YFP (Addgene 101160) and enriched via flow cytometry using dual 646 and YFP select for single clones that contain both 24xMS2 hairpins and stable YFP coat protein expression.

#### Endogenous DNA visualization

WT MCF7 were plated in 24 well at 75% confluency 1 day prior to transfection with 2ul Lipofectamine 2000 (Thermo Fisher) plus 0.3ug DNA_NRIP_gRNA and 0.3ug of NRIP1_Bxb1 suspected in total of 100ul OptiMEM. Cells were incubated in transfection reaction overnight and single clones were screened for Bxb1 landing pad integration. Upon successful clonal validation, selected clone was transfected with 0.3ug pCAGGS_Bxb1o integrase (Addgene 119901) and 0.3ug pDEST_CuOx144_Bxb1 array (Addgene 119903). Single clones were then selected again, PCR validated, and Sanger sequenced for 144xCuO integration. Final to generate stable HaloTag expression corresponding to CuO array, 1ul Super piggyBac Transposase expression vector (System Biosciences PB210PA-1) combined with 0.6ug epB_CymRV5_Halo (Addgene 119907) was transfected according to standard protocol. Stable clones were selected via flow cytometry with Janelia Fluor HaloTag Ligand 646.

#### Endogenous RNA+DNA visualization and ERα/subnuclear structure

Dual RNA+DNA visualization cell line was generated using endogenous DNA stable line as the base. Rather than flow cytometry enrichment using mCherry and HaloTag for endogenous RNA 24xMS2 integration, puromycin selection at 10000x was used instead. Cells after transfection of RNA donor plasmids were selected with puromycin for 3 days. Single clones were then screened for 24xMS2 stable integration after 3 weeks followed by MCP-YFP stable expression.

Endogenous interchromatin granule expression was generated on dual RNA+DNA endogenous line. Post transfection using similar protocol as described above, stable clones where selected via flow cytometry using 3 channels: YFP for RNA, Janelia Fluor HaloTag Ligand 646 for DNA, and mTurquoise2 for SC35.

Endogenous ERα stable line for SMT was generated using similar transfection and selection protocol as previously described. mCherry was used as initial enrichment strategy post 1^st^ round of transfection of gRNA and donor plasmid. Janelia Fluor HaloTag Ligand 646 was used to sort single clones into 96 wells to enrich for ERα HaloTag stable integration.

### Lentiviral transduction

To create virus, 293T wildtype cells were plated at 70% confluency for transfection. Lentiviral pCDH plasmids were transfected along with 2^nd^ generation packaging plasmid psPAX2 (Addgene 12260) and envelop plasmid pMD (Addgene 12259) using lipofectamine 2000. Media was changed the following day and collected at 48 and 72 hours post transfection. Collected viral media was then spun down at 500g for 5 minutes to remove pellet. Viral media was then aliquoted for - 80 °C storage.

Reverse viral transduction was done for viral experiments. Cells were split and seeded for an estimated 20% confluency once sat down in 24 well. Those cells were resuspended and plated in 100% aliquoted viral media plus 1000x polybrene. The viral media was removed the following day and fresh standard culture media was added.

### FISH

#### DNA FISH

MCF7s were plated at 30% confluency on glass-bottom Lab-Tek 8 well chamber slides (Thermo Fisher 177402). Cells were treated with either minus, acute, or chronic E_2_. To induce chronic E_2_ signaling, cells were stripped for 48 hours prior to adding E_2_. To induce acute E_2_ signaling, cells were treated with E_2_ for 1 hour on day 3, prior to fixation protocol. All conditions were fixed with 4% paraformaldehyde on day 3 post stripping for 15 minutes. For DNA FISH, chambers were first washed with 2x SCC twice 3 minutes each. The cells were then treated with 0.1M HCl for 5 minutes at room temperature. HCl was washed away with PBS for 3 minutes, repeated 3 times. Next cells were permeabilized with 0.5% Triton X-100 for 30 minutes at room temperature. The TX-100 was washed away with PBS for 3 minutes, repeated twice. Cells were then incubated in 50% formamide/50% 2x SCC for 1 hour at 37°C. During incubation, DNA FISH probes were prepared, mixing 1ul probe with 4ul hybridizing buffer. Once incubation is done, chamber walls were removed and probe mixture was applied to cells on glass slide and covered with coverslip. Rubber cement was used to seal the edges and slide is protected from light this step forward. The slide is then heated at 80°C for 6 minutes on a heat block and gradually cooled using set program on heat block down to 37°C prior to incubating for 18-24 hours overnight at 37°C in light protected humid chamber. Next day, coverslip is removed and glass slide is washed with warmed 50% formamide/50% 2x SSC wash buffer for 10 minutes, repeated 3 times, at 37°C. The slide is then washed with warmed 2x SSC for 5 minutes, repeated twice, at 37°C. The slide is finally washed once with PBS for 5 minutes and quickly rinsed in distilled water for 3-5 seconds prior to mounting coverslip using Vectashield with DAPI. Coverslip is sealed with nail polish. All *NRIP1* and *TFF1* FISH probes were designed as described in Nair et al. 2019.

Immunofluorescence for ImmunoDNA FISH was performed after 4% PFA fixation but prior to DNA FISH protocol. Fixed cells were first permeabilized with 0.5% Triton X-100 for 20 minutes at room temperature and then washed with PBS. Cells were then blocked in 4% BSA for 1 hour at room temperature. Primary antibodies were diluted to 1:100 in BSA blocking buffer and incubated overnight at 4°C. Primary antibody was washed away the following day using 0.1% PBS-T where 0.1% Triton X-100 was mixed in PBS. Appropriate fluorescent conjugated secondary antibodies were diluted at 1:1000 and cells were incubated for 2 hours. Secondary antibody was washed away with PBS-T prior to continuation with DNA FISH protocol.

#### RNA FISH

MCF7 cells were plated on glass coverslips at 30% confluency. Stripping and E_2_ induction protocol prior to 4% PFA fixation protocol are same as described under DNA FISH. Cells were permeabilized with 70% ethanol and were stored at 4°C if needed. Prior to RNA probe hybridization, coverslips were incubated with Wash Buffer A (2ml Wash Buffer A from Stellaris + 1ml formamide + 7ml water) for 30 minutes at room temperature. RNA FISH probes were resuspended in hybridization buffer (10% formamide + RNA FISH hybridization buffer from Stellaris). Coverslips were incubated with probes overnight at 37°C in a light protected humid chamber. The next day, coverslips were washed once with warmed Wash Buffer A for 30 minutes at 37°C. The second wash is with Wash Buffer A plus Hoechst 3342 diluted at 1:5000 (Thermo Fisher) for 30 minutes at 37°C. The coverslips were then washed with Wash Buffer B (Stellaris) for 5 minutes prior to mounting on glass slide using Vectashield no DAPI. The coverslip is sealed with nail polish. All *NRIP1* and *TFF1* FISH probes were designed as described in Nair et al. 2019.

Immunofluorescence for ImmunoRNA FISH was performed by mixing in primary antibody to RNA FISH probe and then incubated simultaneously overnight. Probes and primary antibody were washed using Wash Buffer A at 37°C as described above followed by fluorescent conjugated secondary antibody incubation of 2 hours prior to second wash with Wash Buffer A plus Hoechst 33342.

#### Validation of RNA and DNA tagged stable lines

Endogenously tagged RNA and DNA stable cell lines were validated via DNA FISH. Live cells were first plated on gridded coverslips (ibidi 10817) at 30% confluency. Stripping and acute E_2_ induction protocol prior to 4% PFA fixation protocol are same as described under DNA FISH. Fixed cells were then imaged immediately on Keyence microscope for punctate signal. After imaging, cells were treated using standard DNA FISH protocol described above. After DNA FISH protocol was completed, those same cells that were imaged following fixation where it maintained fixed live punctate signal were imaged again for FISH signal. The gridded coverslip allowed for accurate identification of location and exact cells previously imaged. Both images were then overlapped according to the grid outline to overlap fixed RNA or DNA endogenous signal with *NRIP1* FISH.

### Microscopy

#### Zeiss LSM 880 Airyscan

Zeiss 880 is equipped with a CO_2_ regulated incubation chamber (Incubator XL S1) and the ambient temperature is held at 37 °C. All imaging was done using the 63x oil objective. All cells were plated on glass-bottom 96 wells (MatTek) prior to live cell imaging. All Zeiss acquired imaging data were Airyscan processed using ZEN software (Zeiss) followed by subsequent analysis described in later individual sections.

For RNA visualization of burst events, endogenous *NRIP1* stably expressing MCP-YFP was imaged using 515 laser in z-stack, definite focus, fast-Airyscan mode at 2-minute intervals. Stripped cells in minus condition were first imaged for 30 minutes prior to addition of E_2_, without disruption in acquisition position. That same patch of cells was continuously imaged in an overnight timelapse immediately after E_2_ treatment to capture acute to chronic transition. For RNA visualization with subnuclear structures of interchromatin granule and Matrin network, the addition of 458 and 594 laser lines were added. To keep the same resolution as single channel RNA visualization, acquisition time interval became 6-minutes.

For DNA visualization of mobility, endogenous *NRIP1* stably expressing CymRV5-HaloTag was imaged using 633 laser in z-stack, definite focus, fast-Airyscan mode at 15-second intervals. Stripped cells in minus condition were first imaged for 10 minutes prior to addition of E_2_, without disruption in acquisition position. That same patch of cells was continuously imaged in an overnight timelapse immediately after E_2_ treatment to capture acute to chronic transition.

For RNA plus DNA dual visualization to measure mobility during burst event, endogenous *NRIP1* stably expressing MCP-YFP and CymRV5-HaloTag was imaged using 515 and 633 lasers combined, in z-stack, definite focus, fast-Airyscan mode at 15 and 30 second intervals. Temporal resolution was maintained with 2-color imaging by zooming into a small patch of cells. Stripped cells were treated with E_2_ and immediately imaged to find patch of cells with burst signal. Once identified, that patch of cells was continuously imaged for 4-6 hours. For RNA, DNA, and interchromatin granule 3-color imaging, the addition of 458 laser was used. Temporal resolution was maintained at 30-second intervals.

DNA and RNA FISH imaging was done using the appropriate laser lines corresponding to fluorescent secondary antibody. Images were taken in z-stack, super resolution Airyscan mode. The only exception is imaging for *NRIP1* endogenous tag FISH overlap validation, which was done on Keyence BZ-X700 microscope using 100x oil objective.

#### Nikon Sterling Eclipse Ti2-E

The Eclipse Ti2-E is equipped with CO_2_ regulated stage top incubator (Tokai Hit) where the temperature is controlled at 37 °C. All SMT experiments were done using the TIRF 100x 1.49 NA objective, with HILO “dirty” TIRF technique. Endogenously Halo tagged ERα cells were treated with Janelia Fluor HaloTag Ligand 549 (Promega GA1110). To prepare HaloTag ligand, a 1:200 dilution was first done to create a 1uM stock. To achieve single molecule tracking, we further diluted the final concentration of HaloTag ligand for imaging to 0.2nM. The cells were stripped and plated on 96 well glass bottom plate as described above. On day of imaging, the cells were incubated in HaloTag Ligand 549 for 20minutes and then washed once. The cells will either be in minus, acute, or chronic E_2_ conditions following the same stimulation protocol as previously described. Once an optimal HILO angle that captures a single plane across the nucleus has been selected, the cells will be excited by 561 laser. Images are acquired every 5ms at 200fps.

For 1,6-hexandiol SMT experiments, cells were treated with either 2%, 3%, 4%, or 5% 1,6-hexandiol (Sigma 240117) dissolved in DMEM white medium. Endogenous HaloTag ERα lines were first imaged as is for baseline. Then the 1,6-hexandiol solution is added for 5 minutes, during which the cells are continuously imaged every minute to monitor change during treatment. The chemical was then washed away with DMEM white medium and replaced with fresh DMEM white medium for recovery. All media followed appropriate protocol as previously described for minus, acute, and chronic E_2_ induction.

### Segmentation of RNA bursts

Maximum projection of RNA bursting images was performed in Zen Black. Subsequent analysis was performed in ImageJ. Nuclei were using the non-focal signal from the RNA, and was detected per frame via Gaussian blurring (sigma=10), background subtraction (rolling ball, rolling=500), autothresholding (Method=mean), default watersheding, and size selection (size=40-200).

Individual bursts were calculated per nucleus per frame in order to avoid edge artefact and spurious burst detection around the nuclear periphery. Individual nuclei were assessed by clearing each frame outside of the periphery of the nuclei being assessed, and then setting the value of the external area to 90% of the median value of the cell. Bursts were then detected per nucleus by DoG (radius 12 and radius 6), and signal was counted as a burst if the DoG levels were above 5+0.55*(cell median intensity). These burst signals were copied into and used to populate an otherwise blank image with values corresponding to the nuclei median, and the process was continued across all nuclei and frames.

### Quantitation of RNA bursts

Segmented bursts were assessed form compilation in a frame-by-frame manner using ImageJ. For each frame, all bursts were compiled using Analyze Particles, and the following values were measured and compiled for each spot: Median nuclei intensity (taken from the value of the spot in the threshold compiled image), Burst intensity (taken from the intensity of the burst area of the original image), frame, and area of burst. These values were compiled, and this process was repeated across all frames. Via the Trackmate plugin for ImageJ, these compiled signals were linked into bursts based on xyzt position, and then quantified to detect the number, frame, and duration of bursts.

### Segmentation of DNA and RNA in DNA-tracking movies

Similarly to how segmentation was done in burst quantitation segmentation, nuclei were first segmented using nuclear signal from background signal in the DNA channel, and segmentation of RNA foci, DNA foci, and organelles were segmented per nuclei and per frame. To reduce memory usage for subsequent processing steps, cells were first manually cropped via ImageJ to into individual cells. Again, per-nucleus analysis allowed for setting on non-nuclear signal to a proportion of the nuclear signal intensity to remove edge artefact, which was applied as before to DNA, RNA, and organelle channels and combined with DoG analysis for segmentation. Organelle segmentation was completed by performing autothresholding of the DoG, DNA thresholding was performed by setting an empirically derived threshold based upon a proportion of the standard deviation of signal in the DNA channel, and RNA thresholding was set to a fixed value throughout images and adjusted upwards or downwards if spurious or insufficient signal was detected.

### Drift correction of segmented DNA-tracking movies

Drift correction was performed on all DNA tracking movies to ensure quantitation of DNA movement within the nucleus free of the motion or rotation of the cell. The Airyscan processed DNA channel was used as the template channel for drift correction, but was first pre-processed to both remove spurious or real focal signal inside the nucleus, as well as to increase the contrast of the nucleus to the extranuclear regions of the image. This was done by first segmenting the target cell nucleus via Gaussian blurring, auto-thresholding (method=Huang2), and an adjustable watershed. The segmented bounds of the cell were dilated and both the nuclear and non-nuclear regions had outlier signal moderated based on median measured values so as to not artefactually skew drift correction. This nuclear-thresholding channel was added to the image being drift correct, and was used via the HyperStackReg for rigid body drift correction. The output was then saved and used for quantitation of DNA motion via Trackmate.

### Quantitation of segmented DNA-tracking movies

Segmented and drift corrected DNA-visualization movies (which may also include segmented RNA and organelle channels) had punctate signal (both RNA and DNA) quantified via the Trackmate plugin for ImageJ. As with RNA burst quantitation, these compiled signals were linked into bursts based on xyzt position, and then quantified to detect the number, frame, and duration of bursts. Distance of DNA signal relative to organelle periphery was determined by subtracting the intensity of the the value of the DNA spot in the internal organelle distance transformed channel from the external value, causing the value for each spot in each track to represent the distance of that spot from the edge of the organelle (positive meaning further outside, negative meaning further inside). RNA burst tracks were used to segment DNA tracks into burst associated or non-burst associated. DNA tracks were thus segmented based on burst or non-burst, or pre-E2, acute (0-120 minutes post E2 treatment), or chronic (post 160 minutes) depending on the experiment and attribute being assessed.

The kinetic properties of DNA tracks were assessed both for the single frame displacement in the segment (e.g. burst vs non burst, or acute versus chronic), as well as mean-squared displacement, diffusion coefficient, and subdiffusivity of segmented tracks. Single frame displacement was calculated by determining the coordinate-based displacement from each frame versus the following frame. Mean squared displacement, diffusion coefficient, and sub diffusivity were calculated in the same mathematical manner as were done in the single molecule tracking experiments above.

### Segmentation of FISH

All FISH image segmentation was performed in ImageJ. Individual nuclei were first identified in the Dapi channel, and were processed first by z-projection, rolling ball background subtraction (rolling=200, sliding window), Gaussian blurring (sigma=4), auto-thresholding using the Huang2 method, removal of small particles (size<20), filling of holes, and a watershed was applied to separate touching nuclei (Adjustable watershed, tolerance =6). Extranuclear signal was cleared for all other channels.

Organelle channel segmentation was performed through Difference of Gaussian (DoG) and autothresholding. The segmented organelle channels and their inversions were then distance transformed using 3D Distance map to generate internal and external distance transformations relative to the edges of these organelles.

Both DNA and RNA FISH channel segmentations were also performed via DoG and autothresholding.

### Quantitation of FISH

Using ImageJ and the 3D Manager plugin, the centroids of segmented FISH dots (both DNA and RNA) were added to the 3D manager. For each, the values of the location of these centroids were assessed in the organelle distance transformed channels (both internal and external) for all organelles assessed. For each FISH dot, the internal distance transformation centroid value was subtracted from the external, providing the distance of each dot from the organelle edge (negative values being inside the structure, positive values being external). These values were then combined over all dots, and compared between conditions and organelles.

### Single Molecule Tracking

#### Pre-tracking processing

Single molecule movies were preprocessed in ImageJ. .Nd2 files of the Red-546 channel and the blue 405 channel were loaded. Auto-thresholding (method=Default) of the single-molecule red channel was performed to determine the area of the nucleus. The average intensity of the red and blue channels were determined over the nucleus area, and these values were recorded in the file names of the image, which was made by saving the red channel as a 32-bit .tif file. The red and blue channel average values were used to confirm similar ERα-Halo expression levels across conditions, while the blue channel levels were used to confirm similar shRNA-expression across conditions.

#### Tracking processing

Tracking of .tif files was performed using the Trackmate plugin in ImageJ. Spot detection was performed using a DoG detector, with a radius of 0.4um and threshold of 10, with no median filtering. Spots before frame 500 were filtered out to select for frames where Halo photobleaching reduces Halo signal to levels where individual molecules are discernable. Tracks were generated by using the Sparse LAP Tracker, with a maximum frame gap of 3, a maximum linking distance of 0.4um, and a maximum closing distance of 0.6um, and a minimum number of spots per track set to 5. All spot statistics were then exported and saved.

#### Calculation of biophysical parameters from tracks

Biophysical properties for tracks were calculated in R. Spots before frame 1000 were filtered out to select for frames where Halo photobleaching reduces Halo signal to levels where individual molecules are discernable. Mean squared displacement was calculated for each track across all dTs that had at least 3 calculable displacements to average. Alpha and diffusion coefficients were calculated using the lm function (linear modeling function, with mean squared displacement ∼ dT) for each track using log-converted mean squared displacement and dT values for all calculated dTs. Radial diffusion coefficient and radium of confinement were similarly calculated from log converted mean squared displacement and dT values using the nls function (MSD ∼ (RADIUSCONF^2) * (1-exp((−4*RADDIFCO*DT)/(RADIUSCONF^2))). The duration of each track remaining within 0.25 and 0.5 um of the starting point of the track were also determined for each track.

#### Clustering

Tracks were clustered using both Gaussian mixture modeling (GMM) and Ward hierarchal clustering in R. To generate a training set for clustering, a randomly selected set of tracks from all wild-type ERα single molecule experiments were taken and balanced evenly across E_2_-, 1h E_2_, and 17h E_2_ conditions. For each track, aforementioned Diffusion Coefficient, Alpha, Radius of Confinement, Radial Diffusion Coefficient, MSD for dT=1, and the duration with a radium of 0.25um were used as dimensions for clustering below.

GMM clustering was performed using the GMM function from the ClusterR library (dist_mode = "maha_dist", seed_mode = "random_subset", km_iter = 20, em_iter = 20, verbose = F) pr = predict_GMM(testdf, gmm$centroids, gmm$covariance_matrices, gmm$weights). The number of clusters used was 6, as this number of clusters resulted in clusters with maximal differences between 1h and 17h E_2_ conditions. This GMM was applied to subsequent SMT experiments to classify tracks as belonging to specific clusters.

Ward clustering was performed using the agnes function from the cluster library, with the “ward” method. The number of clusters used was 6, as this number of clusters resulted in clusters with maximal differences between 1h and 17h E_2_ conditions, which was consistent between this method and the GMM method. A subset of these clustered tracks were then used to train a set of kernalized support vectors using the ksvm function from the kernlab library (scaled = TRUE, type = ‘spoc-svc’, kernel ="laplacedot", kpar = "automatic", C = 8, nu = 0.2, epsilon = 0.1, prob.model = FALSE, class.weights = NULL, cross = 0, fit = TRUE, cache = 40, tol = 0.001, shrinking = TRUE, subset, na.action = na.omi). These support vectors were then applied to subsequent SMT experiments to classify tracks as belonging to specific clusters.

Similar clustering method and algorithm was used on *NRIP1* DNA tracks for all experiments involving flavopiridol treatment.

### RT-qPCR

The cells were stripped for 3 days using phenol red free DMEM with 5% charcoal depleted FBS after reaching 50 – 60% confluency. 100nM E_2_, or ethanol as vehicle, was applied to cells for a time course induction of estrogen signaling including 0, 5, 10, 15, 20, 25, 30, and 60 min. Cells were then collected after treatment and RNA was isolated using Quick-RNA™ Miniprep Plus Kit (Zymo, R1057). 1 μg of total RNA was reverse transcribed by SuperScript III Reverse Transcriptase (Invitrogen, 18080085) followed by qPCR using VeriQuestTM Fast qPCR Master Mix (Thermo Fisher, 75690). Primers used for qPCR are provided. Gene expression levels were normalized based on the expression level of *GAPDH*, and all experiments were conducted with at least two biological replicates and three technical replicates. The following primers were used:

GAPDH_qPCR_F
TGCACCACCAACTGCTTAGC

GAPDH_qPCR_R
GGCATGGACTGTGGTCATGAG

NRIP1_qPCR_F
GCCAAGCCTGTTAGGAACTG

NRIP1_qPCR_R
GGGTGGAGATGGTTTCAAGA

NRIP1e_qPCR_F
CGTCTTTTCCCACTGACACA

NRIP1e_qPCR_R
CCCCTCCCCAGAAGAAAATA

### ChIP-seq

Briefly, approximately 10^7^ cells were cross-linked with 1% formaldehyde in PBS at room temperature for 10 minutes and neutralized with 0.125M glycine. After sonication, ∼50µg soluble chromatin was incubated with 1-5μg of antibody at 4^0^C overnight. Immunoprecipitated complexes were collected using Dynabeads A/G (Invitrogen). Subsequently, immuno-complexes were washed, DNA extracted, and purified by QIAquick Spin columns (Qiagen). For ChIP-seq, the extracted DNA was ligated to Illumina UDI adaptors followed by deep sequencing with Illumina’s HiSeq 4000 according to the manufacturer’s instructions. Sequencing reads were inspected for quality control using FASTQC (https://www.bioinformatics.babraham.ac.uk/projects/fastqc/) and sequencing adaptors were trimmed, if necessary, using TRIMMOMATIC (http://www.usadellab.org/cms/?page=trimmomatic). Reads were aligned to hg19 with Bowtie2(1) (version 2.26) using --very-sensitive setting. Tag directories were then generated using HOMER (2) (version 4.10.3), keeping only unique aligned reads per genome position, allowing one unique read per position (-tbp 1).

### Identification of ChIP-seq Peaks

ChIP-seq peaks were called using HOMER (2) findPeaks subroutine with the default settings (- style factor -o auto). The threshold was set at a false discovery rate (FDR) of 0.001 determined by peak finding using randomized tag positions in a genome with an effective size of 2 × 10^9^ bp. Bedgraph files were generated using HOMER scripts makeUCSCfile and makeMultiWigHub.pl for visualization in the UCSC genome browser. The total number of mappable reads was normalized to 10^7^ for each experiment presented in this study. Heatmaps were generated using deeptools2(3) (version 3.4.3).

### Precision Run-On Sequencing (PRO-seq)

PRO-seq experiments were performed as previously described (4) with modification. Briefly, for nuclei isolation, ∼10 million MCF-7 cells were incubated with swelling buffer (10 mM Tris-HCl pH7.5, 2 mM MgCl_2_, 3 mM CaCl_2_) for 5 minutes on ice and then incubated with lysis buffer (swelling buffer with 0.5% NP-40 and 10% glycerol) for 5 minutes on ice, before being re-suspended in 100μl of freezing buffer (50 mM Tris-HCl pH8.0, 40% glycerol, 5 mM MgCl_2_, 0.1 mM EDTA). For Run-on assay, an equal volume of reaction buffer (10 mM Tris-HCl pH 8.0, 5 mM MgCl_2_, 300 mM KCl, 1 mM DTT, 20 units of SuperaseIn, 1% sarkosyl, 500 μM ATP, GTP, bio-UTP, and bio-CTP) was added into each sample before incubation at 30°C for 5 min. The nuclear run-on RNA was then extracted with TRIzol LS reagent (Invitrogen) and subjected to hydrolysis, buffer exchange and purification by streptavidin beads (Thermo Fisher). Purified RNA was treated with PNK before being used for cDNA synthesis by using NEBNext® Multiplex Small RNA Library Prep Set for Illumina® Kit (NEB). Obtained cDNA templates were amplified by PCR using the LongAmp® Taq 2X Master Mix (NEB) for deep sequencing.

For PRO-seq experiments, sequencing reads were inspected for quality control using FASTQC (https://www.bioinformatics.babraham.ac.uk/projects/fastqc/) and sequencing adaptors were trimmed, if necessary, using TRIMMOMATIC (http://www.usadellab.org/cms/?page=trimmomatic). Reads were aligned to hg19 with Bowtie2 (1) (version 2.26) using --very-sensitive setting. Tag directories were then generated using HOMER (2) (version 4.10.3), keeping only unique aligned reads per genome position, but allowing up to 3 unique reads per position (-tbp 3). For read counting, the HOMER script analyzeRepeats.pl was used to estimate the raw counts per gene. The aligned reads were counted over the RefSeq gene bodies. Bedgraph files were generated using HOMER scripts makeUCSCfile and makeMultiWigHub.pl for visualization in the UCSC genome browser. Heatmaps were generated using deeptools2(3) (version 3.4.3).

#### Data Availability

ChIP-seq and PRO-seq data that support the findings of this study have been deposited in GEO with the accession code <u>GSE221899</u>. Please enter token <u>wvibgywqjpuzzwf</u> to access.

#### Code Availability

Code used for all imaging data related analysis and graphs has been uploaded and available at: https://github.com/susanxwang/Imaging.

